# inMTSCCA: An Integrated Multi-task Sparse Canonical Correlation Analysis for Multi-omics Brain Imaging Genetics

**DOI:** 10.1101/2022.10.30.514398

**Authors:** Lei Du, Jin Zhang, Ying Zhao, Muheng Shang, Lei Guo, Junwei Han, the Alzheimer’s Disease Neuroimaging Initiative

**Affiliations:** Department of Intelligent Science and Technology, School of Automation, Northwestern Polytechnical University, Xi’an 710072, China

**Keywords:** Brain imaging genetics, multi-omics endophenotypes, cross-endophenotype association, genetic risk factors

## Abstract

Identifying genetic risk factors for Alzheimer’s disease (AD) is an important research topic. To date, different endophenotypes such as imaging-derived endophenotypes and proteomic expression-derived endophenotypes have shown the great value in uncovering risk genes compared to case-control studies. Biologically, a co-varying pattern of these different omics derived endophenotypes could result from the shared genetic basis. However, existing methods mainly focus on the effect of endophenotypes, and that of cross-endophenotype associations remains largely unexploited. In this paper, we used both endophenotypes and their cross-associations of multi-omics to identify genetic risk factors, and proposed two integrated multi-task sparse canonical correlation analysis (MTSCCA) methods, i.e., pairwise endophenotype correlation guided MTSCCA (*pc*MTSCCA) and high-order endophenotype correlation guided MTSCCA (*hoc*MTSCCA). *pc*MTSCCA employed pairwise correlations between MRI-derived, plasma-derived, and cerebrospinal fluid (CSF) derived endophenotypes as an additional penalty. *hoc*MTSCCA used high-order correlations among these multi-omics for regularization. To figure out genetic risk factors at individual and group levels, as well as altered endophenotypic markers, we introduced sparsity-inducing penalties in both models. We compared *pc*MTSCCA and *hoc*MTSCCA with three related methods on both simulation data and real neuroimaging, proteomic analytes, and genetic data. The results showed that our methods obtained better or comparable canonical correlation coefficients (CCCs) and feature subsets than benchmarks. Most importantly, the identified genetic loci and heterogeneous endophenotypic markers showed high relevance. Therefore, jointly using multi-omics endophenotypes and their cross-endophenotype associations are promising to reveal genetic risk factors, and both methods are qualified for this complicated task. The source code and manual of inMTSCCA is available at: https://ngdc.cncb.ac.cn/biocode/tools/BT007330.

## Introduction

Alzheimer’s disease (AD) is one of the severe brain disorders and has been known as highly inheritable [1]. AD usually attack many components of the body system, *i*.*e*., the brain tissue, the blood system and the cerebrospinal fluid (CSF) [2], which could lead to many abnormal alterations. Therefore, AD patients could manifest with multiple altered endophenotypes, *i*.*e*., the measurable traits at different levels of biological organization. These alterations involve the magnetic resonance imaging (MRI) derived quantitative traits (QTs) and plasma-derived proteomic analytes, could co-occur without coincidence. The co-occurrence of multiple heterogeneous endophenotypes could play a critical role in identifying genetic risk factors of AD, given the success of multiple endophenotypic traits genetic association studies [3-5]. The co-occurrence of multiple endophenotypes could result in a relative high correlation between multiple endophenotypes and is quantified as the cross-endophenotype (CEP) association in this paper. Taking AD as an example, the apolipoprotein E(APOE) genotype is synchronously associated with different endophenotypes, including APOE protein levels in the cerebrospinal fluid (CSF) [6-8], as well as neuroimaging QTs of the reduced hippocampus volume and elevated amyloid deposition [9]. That is, both CSF levels and imaging QTs could point to the same genetic basis, and thus they may be correlated in the absence of AD [10]. On this account, the CEP association, in this paper, exists because a genetic locus or gene is associated with more than one endophenotypes regardless of the underlying cause [3]. Since the CEP association can occur within omics data (e.g., CEP associations within imaging QTs), and between multi-omics (e.g., CEP associations between imaging QTs and proteomic analytes), both intra- and inter-omics CEP associations could point to shared genetic factors. Therefore, the multi-omics CEP association stands a good chance of prompting the identification of genetic risk factors, which would yield new insight into the genetic architecture of AD. Pleiotropy is a similar terminology, which takes the causal effect into consideration. It refers to that a genetic locus or gene truly affects multiple endophenotypes [3]. Therefore, the pleiotropy generally focuses on identifying causal variants other than tag SNPs. With the aim of detecting comprehensive genetic factors, the CEP association is a better choice since it has more comprehensive coverage of genetic effects. Generally, we can further distinguish between different types of pleiotropy (such as biological pleiotropy, mediated pleiotropy, and spurious pleiotropy) based on the output yielded by multi-omics CEP association studies [3, 11].

In brain imaging genetics, different endophenotypes have been widely used where the associations between endophenotypes, e.g., imaging QTs or proteomic analytes, and single nucleotide polymorphisms (SNPs) were extensively investigated [12]. Compelling evidence suggests that using carefully selected endophenotypes, e.g., AD-altered imaging QTs, is able to discover novel loci that are unlikely to be revealed in case-control studies [12,13]. Roughly speaking, there are three distinct types of analytical methods, including the univariate method, the multivariate regression method and the bi-multivariate correlation method [12].

In brief, the univariate methods repeatedly analyze every pair of QT and SNP [9]. The multivariate regression methods investigate the impact of a segment of SNPs on one or multiple QTs, which can capture the group structure of multiple SNPs simultaneously [14]. The bi-multivariate correlation methods study the association between multiple QTs and multiple SNPs (multi-QTs-multi-SNPs). To the best of our knowledge, a common critical issue of these methods is that they either ignore the CEP association [9], or only involve the intra-omics CEP one [15-17]. As a result, the inter-omics CEP association remains largely unemployed. Therefore, it is essential and important to develop new methods that can employ multi-omics CEP associations including both intra- and inter-omics ones, which could gain increased capability in identifying meaningful and reliable genetic loci [11].

Although the multiple traits genetic association study could be an effective tool for this task. They still face two difficult challenges. First, they usually use peripheral phenotypic traits which could have limited identification power compared to endophenotypic traits [3-5]. Second, they emphasize on a limited set of pre-selected traits which may ignore the associations of trait pairs, especially those seemingly distinct trait pairs being synergistically regulated by shared genetic basis [3].

In this paper, to overcome the above drawbacks, we propose two integrated multi-task sparse canonical correlation analysis (MTSCCA) methods to identify genetic factors. These two MTSCCA methods, called the pairwise endophenotype correlation guided MTSCCA (*pc*MTSCCA) and the high-order endophenotype correlation guided MTSCCA (*hoc*MTSCCA), incorporate different types of CEP associations. *pc*MTSCCA utilizes the pairwise CEP associations between MRI-derived, plasma-derived, and cerebrospinal fluid (CSF) derived endophenotypes as an additional penalty. *hoc*MTSCCA incorporates the high-order CEP associations among MRI-derived, plasma-derived and CSF-derived endophenotypes for regularization. To identify meaningful genetic risk factors and relevant endophenotypes of multi-omics, we use L_21_-norm, L_1_-norm and a newly designed Fused pairwise Group Lasso (FGL_21_) penalty in conducting feature selection at different levels. The contributions of this paper are fourfold. First, we employ both multi-omics endophenotypes and their CEP associations, which is a better modelling strategy than existing methods. Therefore, both *pc*MTSCCA and *hoc*MTSCCA possess a comprehensive and accurate ability for risk loci identification. Second, both pairwise and high-order associations are used and verified, which could provide a diverse and useful guidance for future method development. Third, the combination of L_21_-norm and FGL_21_ aid to select the shared risk loci affecting multi-omics data jointly. The FGL_21_ penalty further takes the linkage disequilibrium (LD) into consideration, which is a practical feature selection penalty. Fourth, we propose a unified and efficient iteration optimization algorithm, and theoretically analyzed its convergence.

To evaluate both *pc*MTSCCA and *hoc*MTSCCA, we compare them with three related methods, i.e. one conventional sparse multiple CCA (SMCCA) [18] and two state of-the-art ones (Adaptive SMCCA [19] and RelPMDCCA [20]), since they are suitable for multi-omics data. We use four simulation datasets with different weight patterns and signal-to-noise (SNR) levels, and one real application dataset containing the brain imaging QTs, proteomic analytes and SNPs from Alzheimer’s disease neuroimaging initiative (ADNI) [21]. The goal is to employ both multi-omics endophenotypes and their CEP associations to identify a comprehensive and meaningful subset of AD-risk loci, as well as those relevant heterogeneous endophenotypes including imaging QTs and proteomic analytes. In the simulation study, both *pc*MTSCCA and *hoc*MTSCCA obtain higher or comparable canonical correlation coefficients (CCCs) and better canonical weight profiles than all competing methods. In the real study, *pc*MTSCCA and *hoc*MTSCCA again outperform all competitors in terms of higher CCCs and cleaner canonical weight patterns. By looking into the identified biomarkers, we find that most of the identified imaging QTs, proteomic analytes and SNPs are related to AD individuals. In contrast, the competitors yield too many signals with both relevant and irrelevant biomarkers reported, which could mislead the subsequent further analysis. To summarize, both integrated MTSCCA methods are good at fusing multi-omics data to detect interesting biomarkers, which would offer a very promising new strategy for brain imaging genetics and multi-omics studies, and further deepen our understanding of the etiology and pathology of AD.

## Results

### Experimental setup

To evaluate the proposed methods, we chose three related methods which can analyze the associations among multi-omics data and select relevant feature subsets simultaneously. They were the conventional L_1_-norm penalized SMCCA [18] and two state-of-the-art ones such as L_1_-norm penalized Adaptive SMCCA [19] and SCAD penalized SMCCA (RelPMDCCA) [20]). Using both conventional and state-of-the-art methods can make valid and thorough comparisons. Those other SCCA methods were not included because they cannot handle multi-omics data directly.

We employed the nested five-fold cross-validation strategy fine-tuned parameters. In the inner loop, every combination of the candidate parameters was evaluated by cross-validation, and those yielding the best testing values were optimal parameters. Receiving the optimal parameters from the inner loop, the outer loop was used to generated final training and testing results. In this paper, we set the range of candidate parameters to [0.01, 0.1, 1, 10, 100]. In some cases, this procedure might not find desirable parameters. The reason is that we usually have two goals at the same time. One is to identify a high correlation, and the other is to select relevant features. During the parameter tuning, we only focused on the correlation coefficients. To overcome this, we employed a two-stage strategy. The first stage used a large interval as we just described. In the second stage, we used a small interval to further seek the optimal parameters. In particular, if Θ was the parameters obtained from the first stage, the second stage will tune in Θ±[0.1, 0.2, …, 0.5]. In practice, this setup usually yielded reasonable parameters for both correlation and feature selection. In the experiments, all methods ran on the same setup such as the same data partition, candidate parameters and software platform, which can make the comparison fair.

In this study, we employed the Tensor Toolbox software (https://gitlab.com/tensors/tensortoolbox) [25] for *hoc*MTSCCA. In addition, we used feature selection ability and canonical correlation coefficient (CCC) as indicators to evaluate the performance of the model. The CCC can be calculated as follows

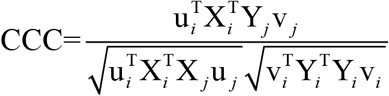

when both X and Y_i_’s are normalized.

### Simulation study

We generated four simulated datasets with different ground truths to ensure a thorough comparison. Each dataset had four omics data to simulate SNPs, plasma proteins, CSF proteins, and imaging QTs. Data 1 and Data 2 were generated from the same ground truth, but with different signal to noise (SNR). Data 3 was different to the previous two ones with different positions and directionality of true signals. Data 4 had a distinct number of feature dimensionality to simulate a small-*n*-large-*p* problem. We also designed group structures to simulate the LD of genetic data for these datasets. The ground truth of each dataset was shown below and in **Figure 1** (top row).

**Figure 1.** Canonical weights on synthetic data from 20 times five-fold cross-validation. Each row is a method: (1) Ground truth; (2) SMCCA; (3) Adaptive SMCCA; (4) RelPMDCCA; (5) *pc*MTSCCA; and (6) *hoc*MTSCCA. Each column corresponds to a canonical weight. The first column u is the canonical weight of X, the second column v_1_ is that of Y_1_, the third one is the canonical weight of Y_2_, and the last column is that for Y_3_. The values in each panel are obtained from 20 times trials.

Data 1: 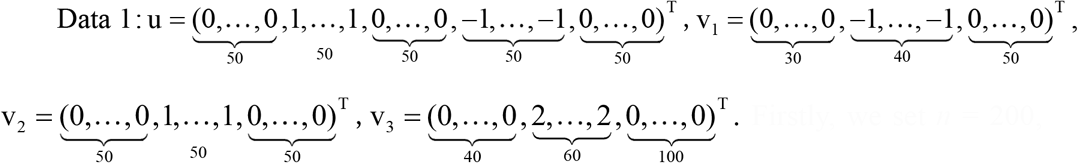. Firstly, we set *n* = 200, *d* = 250, *p*_1_ = 120, *p*_2_ = 150, *q* = 200, where *n* denoted the number of subjects, *d* was the number of SNPs, *p*_1_ was that of plasma-derived proteomic markers, *p*_2_ was the number of CSF-derived proteomic markers, and *q* was the number of imaging QTs. Then we generated four sparse vectors, *i*.*e*., u ∈ ℝ ^*d*×1^, and three sparse vectors 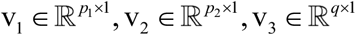. Using a latent vector z ∼ *N*(0,*σ* × I) with *σ* = 0.05, we created X by x^*l*^ ∼ *N* (z_*l*_ u^T^, *σ* ×∑_*x*_)|, where x^*l*^ was the *l* − *th* row of X, and 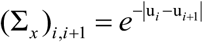 simulated the group structures of SNPs. Finally, we created each Y_*t*_ by 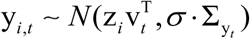, where 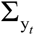 was an identity matrix. The details of the data generation procedure can be found in [26].

Data 2: This data set used the same settings as the first one, but with different noise levels, *i*.*e*., *σ* = 0.2.

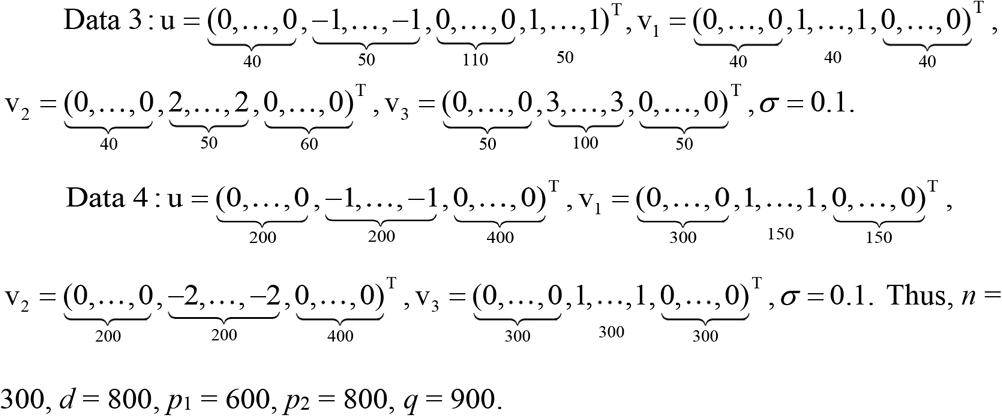

We applied all methods to four simulated datasets. We employed the canonical correlation coefficients (CCCs) and canonical weight patterns to evaluate each method. A higher CCC stands for a better performance. **Table 1** presented the averaged CCCs from 20 times five-fold cross-validation. For ease of presentation, we denoted the CCC between X and Y_1_ as CCC1-1, that between X and Y_2_ as CCC1-2 and so on. We observed that both *pc*MTSCCA and *hoc*MTSCCA obtained higher or comparable CCCs than benchmark methods on both training and testing sets for most cases. In particular, the first two datasets were generated with the same ground truth but different noise levels, and results on Data 1 and Data 2 demonstrated that our methods can outperform benchmarks under different intensities of noise, indicating a robust performance. In addition, both proposed methods also yielded better CCCs than benchmarks on Data 3 and Data 4 where the true signals and number of features were different. In most cases on the testing CCCs, RelPMDCCA held the largest standard deviations attributing to its unstable performance caused by the SCAD penalty.

Besides CCCs, the heatmaps showing the canonical weight patterns were shown in **Figure 1**. Each benchmark had only one canonical weight u, and we repeatedly showed them for three times. This enables a clear comparison between all methods. In each panel, the average weight from 20 times trials was shown. From the figure we observed that RelPMDCCA, *pc*MTSCCA and *hoc*MTSCCA identified a small feature subset which were consistent to true signals. On the contrary, SMCCA and Adaptive SMCCA cannot identify where the true signals were. This could due to their independent assumption which might miss important information. In summary, these simulation results demonstrated that by eliminating the independent assumption, a method could be good at feature selection. Additionally, by fusing both multi-view data and their cross-associations, a better CCC could be obtained. We will verify this conclusion on the real ADNI data next.

### Real neuroimaging, proteomic, and genetic study

The brain imaging data, quantification of proteomic analytes in plasma and CSF, and genotyping data were obtained from the ADNI database. The primary goal of this initiative is to test whether serial MRI, or other biological markers, and clinical and neuropsychological assessment can be combined to measure the progression of mild cognitive impairment (MCI) and early AD. For up-to-date information, see www.adni-info.org.

There were 244 non-Hispanic Caucasian participants containing 42 healthy controls (HCs), 137 MCIs and 65 Ads, and the details of the participant characteristics are shown in **Table 2**. Their baseline structural MRI scans were collected and pre-processed via a widely used pipeline including the average, alignment, resample, smoothness and normalization steps. To ensure the efficiency and increase the power, we extracted the region of interest (ROI) level voxel-based morphometry (VBM) in the SPM software. Finally, we obtained 465 imaging QTs which were ROI measurements (grey matter density measures) spanning the whole brain based on the MarsBaR anatomical automatic labelling (AAL) atlas [27]. These imaging QTs were also adjusted to eliminate the effects of the baseline age, gender, handedness and years of education.

The blood plasma and CSF samples of these 244 subjects were evaluated by Rules Based Medicine, Inc. (RBM) proteomic panel. After quality control (QC), we separately generated 146 plasma-derived proteomic markers and 83 CSF-derived ones. These subjects’ genotyping data were genotyped using the Human 610-Quad or OmniExpress Array platform (Illumina, Inc., San Diego, CA, USA). These genotype data were then pre-processed according to the standard quality control (QC) and imputation steps. We finally contained 1,094 SNPs located in the neighbour of AD risk gene *APOE* (boundary: ± 200kb) according to the ANNOVAR annotation. We aim to identify AD-risk loci by these heterogeneous multi-omics endophenotypes, AD-affected endophenotype markers, as well as the bi-multivariate association between imaging QTs, proteomic markers and SNPs.

In this real ADNI data, we presented the CCCs of all methods in **Table 3** for comparison. We denoted the CCC between SNPs and plasma-derived proteomic markers as SNP-Plasma for simplicity. Those remaining CCCs can be represented similarly. We observed that both *pc*MTSCCA and *hoc*MTSCCA obtained higher CCCs, including both training and testing scores, for most cases than comparison methods. In particular, our methods obtained much higher CCCs than competitors for SNP-plasma and SNP-CSF CCCs. They also outperformed competitors for the SNP-VBM CCCs on testing sets. This implies that both *pc*MTSCCA and *hoc*MTSCCA yielded higher values between SNPs and proteomic markers than those between SNPs and brain imaging QTs of structural MRI scans. This is very interesting since it is in agreement with the biological organization. But all comparison methods failed to verify this. This is also the evidence suggesting that using both multi-omics and their CEP associations can reasonably capture the relationship among multiple distinct omics data in real studies.

### Identification and interpretation of genetic loci

The heatmap in **Figure 2** showed the canonical weights corresponding to the genetic data for each method. We clearly observed that *pc*MTSCCA and *hoc*MTSCCA identified multiple relevant AD-risk loci, including the notorious common variant rs429358 (*APOE*) [28], rs56131196 (*APOC1*), rs4420638 (*APOC1*), rs7412 (*APOE*), rs440446 (*APOE*) and so forth. In **Table 4**, all the top ten genetic loci of *pc*MTSCCA and *hoc*MTSCCA have been shown to increase the risk of AD. Although SMCCA and Adaptive SMCCA could also identify some AD-related loci, they reported too many signals which indicates that they consider most of SNPs are related to AD. Obviously, this is unconscionable. RelPMDCCA yielded a sparser result than SMCCA and Adaptive SMCCA but still denser than ours. And, it missed the strongest rs429358 which makes its identification unconvincing. In addition, *pc*MTSCCA and *hoc*MTSCCA clearly identified two group of SNPs, which was supported by their newly designed FGL_2,1_ penalty. Specifically, SNPs with equal or very similar weight values will be reported in a same group. More details can be observed in **Table S1** in the supplementary. After investigation, we found that SNPs in the same group were truly from the same LD, which demonstrated that our methods were better than competitors in terms of both individual level and structural level feature selection.

**Figure 2.** Canonical weights (mean value) of SNPs from 20 times fivefold cross-validation. Each row is a method: (1) SMCCA; (2) Adaptive SMCCA; (3) RelPMDCCA; (4) *pc*MTSCCA; and (5) *hoc*MTSCCA.

### Identification and interpretation of plasma-derived proteomic markers

The canonical weights showing the importance of plasma-derived proteomic markers were shown in **Figure 3**. The figure clearly exhibited that *pc*MTSCCA and *hoc*MTSCCA identified a small subset of all plasma-based markers by assigning them a relative higher weight value. These identified markers, such as ApoE, ApoB, ApoC-I, CRP, CD5L, CgA, MIG, Testosterone and so on [29-32], have been verified to be related to AD. In contrast, SMCCA and Adaptive SMCCA failed to convey useful information for the excessively reported signals. Due to the SCAD penalty, RelPMDCCA obtained too sparse signals with many risk plasma markers missed. To make the comparison clear, we also presented the top ten identified markers in **Table 5**, and the table with more details such as the weight values was contained in the supplementary (**Table S2**). Next, we will show the results in terms of CSF-derived proteomic markers.

**Figure 3.** Canonical weights (mean value) of plasma-derived proteomic markers from 20 times five-fold cross-validation. Each row is a method: (1) SMCCA; (2) Adaptive SMCCA; (3) RelPMDCCA; (4) *pc*MTSCCA; and (5) *hoc*MTSCCA.

### Identification and interpretation of CSF-derived proteomic markers

Figure 4 showed heatmap regarding canonical weights of CSF-derived proteomic markers, and **Table 6** contained the top ten selected CSF markers (more details were presented in **Table S3**). It is interesting that both our methods successfully identified AD-related CSF-derived proteomic markers including ApoE, CRP, FABP-heart, MIG and so forth [33, 34]. Both methods identified the FGF-4 as the most relevant and further investigation should be warranted. SMCCA and Adaptive SMCCA again yielded too many markers which were very hard to interpret. Thus, they both could hardly convey useful information in real studies. RelPMDCCA committed the same mistake as it did for plasma-derived markers identification. Its over-sparse results are prone to miss some meaningful markers such as ApoE. Combining results obtained from both plasma- and CSF-derived data, we concluded that both *pc*MTSCCA and *hoc*MTSCCA performed much better than benchmarks, demonstrating their superior performance in identifying relevant proteomic expression markers.

**Figure 4.** Canonical weights (mean value) of CSF-derived proteomic markers from 20 times five-fold cross-validation. Each row is a method: (1) SMCCA; (2) Adaptive SMCCA; (3) RelPMDCCA; (4) *pc*MTSCCA; and (5) *hoc*MTSCCA.

### Identification and interpretation of imaging QTs

Identifying AD-affected brain regions is also important to characterize AD. In **Figure 5**, canonical weights corresponding imaging QTs were shown. We also presented the top ten selected imaging QTs in **Table 7** (see **Table S4** in the supplementary for more details) to make a clear comparison. We observed that for this structural MRI data, *pc*MTSCCA and *hoc*MTSCCA reported that both left and right hippocampus were the most areas being affected by AD. The left middle frontal cortical and left parahippocampal gyrus were also vulnerable. This was in agreement to previous studies that severe atrophy happened to these areas in AD patients. The benchmarks could not provide helpful information since they reported too many irrelevant imaging QTs. That is, they considered that almost all brain areas were vulnerable to AD, which was meaningless for a clinician since he/she cannot easily select the most relevant imaging QTs for further investigation. To sum up, results on SNPs, two types of proteomic makers and imaging QTs jointly demonstrated that, *pc*MTSCCA and *hoc*MTSCCA can not only identify meaningful genetic loci at individual level and LD level, but also detect heterogeneous AD-related endophenotypes including plasma- and CSF-derived proteomic makers and imaging QTs. This further demonstrated that jointly using multi-omics endophenotypes and their CEP associations could be a promising direction to accurately and comprehensively identify genetic risk factors, as well as relevant heterogeneous endophenotypes they underpin. All these conclusions confirmed that our methods are powerful and practical for multi-omics fusion analysis, which can yield important clues for subsequent analysis.

**Figure 5.** Canonical weights (mean value) of brain imaging QTs from 20 times five-fold cross-validation. Each row is a method: (1) SMCCA; (2) Adaptive SMCCA; (3) RelPMDCCA; (4) *pc*MTSCCA; and (5) *hoc*MTSCCA.

### Refined analysis of identified biomarkers

We now investigate the association between SNPs and plasma-derived proteomic biomarkers, CSF-derived ones, as well as imaging QTs. The ANOVA was also conducted to verify the effectiveness of the CEP association in identifying risk loci.

### Association between selected SNPs and the individual endophenotype

The pairwise associations between SNPs and three separate types of endophenotypes, including the plasma-derived proteomic expression biomarkers, CSF-derived ones and imaging QTs were shown in **Figure 6** and **Figure 7** for *pc*MTSCCA and *hoc*MTSCCA respectively. Those significant values (*p* < 0.05) were marked by “×” symbols. It was clear that most association scores reached the significant level, indicating the effectiveness of these markers identified by our methods. In **Figure 6**, the subfigure (A) showed that the apolipoprotein E (ApoE) concentration level was significantly correlated to nine out of ten SNPs, showing its high relationship to these identified AD-risk loci. In addition, rs429358, rs56131196 and rs4420638 shared the same correlation patterns to proteomic markers herein, which implicated that these three loci were located in the same group. A literature search confirmed that these three loci were in the same LD block, which demonstrated the merit of the FGL penalty. The same patterns can be observed in **Figure 7** too. Both figures showed that our methods could figure out meaningful pairwise association between SNPs and endophenotypes.

**Figure 6.** *pc*MTSCCA’s pairwise correlation. (a) The association between identified SNPs and plasma-derived proteomic biomarkers. (b) The association between identified SNPs and CSF-derived proteomic biomarkers. (c) The association between identified SNPs and imaging QTs. The ‘×’ symbol indicated that this pairwise association reached the significance level (p < 0.05).

**Figure 7.** *hoc*MTSCCA’s pairwise correlation. (a) The association between identified SNPs and plasma-derived proteomic biomarkers. (b) The association between identified SNPs and CSF-derived proteomic biomarkers. (c) The association between identified SNPs and imaging QTs. The ‘×’ symbol indicated that this pairwise association reached the significance level (p < 0.05).

### Association between selected SNPs and the pairwise cross-endophenotype association

To show the effectiveness of the pairwise cross-endophenotypes association, we conducted ANOVA to investigate the effects of two (plasma- and CSF-derived) protein expression levels, imaging QTs and their two-way interactions (associations) on SNPs with age, gender, years of education and handedness as covariates. We focused on investigating rs7412 which was missed by those comparison methods without consideration of the CEP association. The main effects of the concentration of plasma-APOE and CSF-APOE were all significant (*p* = 5.26×10 ^−7^ and *p* = 4.31×10 ^−5^), but that of hippocampal volume was not (*p* = 0.33). The interesting and meaningful things are that the pairwise association between plasma-APOE and CSF-APOE concentrations, and that between plasma-APOE concentrations and hippocampal volume were significant. This indicated that the pairwise association between heterogeneous endophenotypes could be beneficial to risk loci identification.

### Association between selected SNPs and the high-order cross-endophenotype association

The effects of high-order CEP association in assisting identification of risk loci are also of interest. The age, gender, years of education and handedness were included as covariates. The ANOVA results showed that main effects of both plasma-APOE and CSF-APOE were significant with *p* = 5.92×10 ^−7^ and *p* = 4.68×10 ^−5^ respectively. Surprisingly, the effect of high-order CEP among plasma-APOE, CSF-APOE and hippocampal volume reached the significant level (*p* = 4.95×10 ^−4^) even though the hippocampal volume alone was insignificant.

Combined results here and that of the last subsection, we concluded that both pairwise and high-order CEP associations could implicate risk genetic loci, which, in all probability, is due to the shared genetic mechanism of these multiple endophenotypes. Therefore, both proposed computational methods are qualified for brain imaging genetics by taking the CEP association into consideration.

### Comparisons of identified biomarkers among different diagnostic groups

In this subsection, we investigated that if the selected phenotypic markers were distinctly distributed among different genotypes and diagnostic groups such as HC, MCI and AD. **Figure 8** showed the phenotypic distributions for plasma (APOE), CSF (APOE) and hippocampal volume. In sub-figure (A), decreased plasma’s (APOE) concentration level was observed in both MCI and AD groups, and the concentration level of ADs was lower than that of MCIs. Within each diagnostic group, the homozygote TT and heterozygote CT showed decreased Alzheimer’s risk, indicating that the minor allele T could be an AD-inhibited allele since holding this nucleotide exhibited a low risk of AD. On the contrary, carrying the major allele C might be an AD-risk factor as these participants were more vulnerable to AD attack. Similar conclusions could also be observed in sub-figures (B) and (C). The brain atrophy in sub-figure (C) showed that the nucleotide C showed increased risk to ADs, but this could not be observed in homozygous CC in HC group. These results demonstrated that the concentration level of plasma (APOE), CSF (APOE) and hippocampal volume could be indicators for AD diagnosis, and the allele C of rs7412 might be an implicit factor for developing AD.

**Figure 8.** Pairwise comparisons for SNP and endophenotype among different diagnostic groups such as HC, MCI and AD respectively. (a) The concentration of plasma-APOE for different genotypes among three groups. (b) The concentration of CSF-APOE for different genotypes among three groups. (c) The atrophy in the left hippocampus lobe for different genotypes among three groups.

To summarize, the statistical analysis confirmed the value of the CEP association in assisting the identification of risk loci. In addition to the endophenotype itself, their correlation could be a good measurement in figuring our meaningful risk loci, which could increase our understanding of the genetic mechanism of AD.

## Discussion

We have showed that both the pairwise CEP association and high-order one can improve the bi-multivariate association and feature selection. *pc*MTSCCA employs multi-omics endophenotypes and their pairwise CEP association with due consideration given to the computation complexity since the pairwise correlation is easy to calculate. *hoc*MTSCCA uses multi-omics endophenotypes and the high-order CEP association, but spends more time than *pc*MTSCCA due to the tensor calculation. As a result, we suggest using *pc*MTSCCA if one intends to obtain the results quickly. On the other hand, *hoc*MTSCCA is a good alternative if one prefers the biomarker identification to time consumption.

## Materials and methods

In this article, we denote vectors as lowercase letters, and matrices as uppercase letters. The *i*-th row and *j*-th column of X=(*x*_*ij*_) are denoted as x^*i*^ and x_*j*_ respectively. Besides, the Euclidean norm of x is defined as 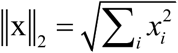,The L_21_-norm of X is defined as ∥X∥_2,1_ = ∑_*i*_∥*x*^*i*^∥_2_,and the Frobenius norm is defined as 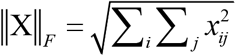

### Background

In this subsection, we first briefly introduced the related sparse multiple canonical correlation analysis (SMCCA), the Adaptive SMCCA and the Relaxed PMDCCA (RelPMDCCA).

#### Sparse multiple canonical correlation analysis (SMCCA)

SMCCA is an extension of the two-view SCCA which can mine the associations among more than three views. Suppose we face a multi-omics issue where the imaging QTs, proteomic expression markers and SNPs are provided, SMCCA is suitable to calculate their complex associations. For ease of presentation, SNPs, plasma-derived proteomic markers, CSF-derived ones and imaging QTs are associated with 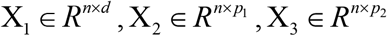 and X4 ∈ *R*^*n*×*q*^ respectively, where *n* is the number of subjects, *d* is the number of SNPs, *p*_1_ is the number of plasma-derived markers, *p*_2_ is the number of CSF-derived markers, and *q* is the number of imaging QTs. Now the SMCCA can be formulated as follows

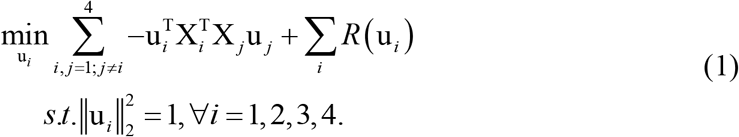

Here u_*i*_’s are the canonical weights for four omics data which indicates the contribution of each biomarker. R(u_*i*_)’s are the regularization terms (e.g., L_1_-norm) to figure out a small subset of biomarkers with the highest relevance. Generally, there are tuning parameters to balance between the loss function and regularization terms.

According to [22], SMCCA requires the SNP data to be associated with multiple heterogeneous endophenotypes simultaneously. This is overstrict and thus could be suboptimal since loci affecting one omics data alone might be missed. Another obvious shortcoming is that SMCCA takes the independent assumption, *i*.*e*., 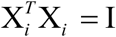, which deems all features are independent. This could pay for the performance degradation [20].

#### Adaptive SMCCA

The Adaptive SMCCA improves SMCCA via adding an additional tuning parameter. Formally, Adaptive SMCCA is given as follows

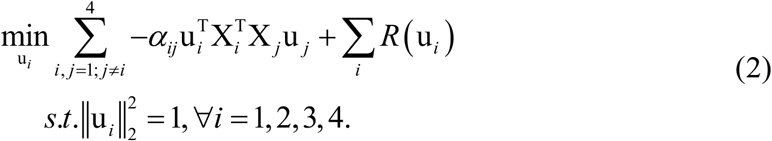

*αij*^’^*s* are parameters to balance between different sub-objectives since the independent assumption of 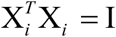 will lead to biased optimization [19]. In general, Adaptive SMCCA is similar to SMCCA. For one thing, it still depends on the independent assumption. For another, it is too strict to require SNPs to be associated with multi-omics data simultaneously, which is suboptimal.

#### RelPMDCCA

By now, RelPMDCCA is the latest and best SMCCA which has been proposed to analyze multi-omics data simultaneously [20]. The definition of RelPMDCCA is similar to SMCCA, i.e.

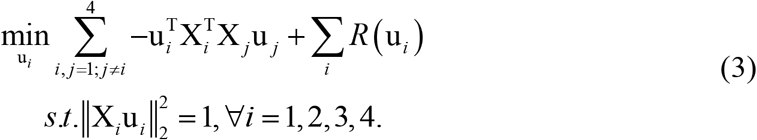

However, RelPMDCCA is different to SMCCA in two aspects. First, it gets rid of the independent assumption. Second, RelPMDCCA employs the smoothly clipped absolute deviation (SCAD) penalty, which could be a more appropriate surrogate of the ideal L_0_-norm than SMCCA’s L_1_-norm.

In summary, all above three SMCCA methods, including SMCCA, Adaptive SMCCA and RelPMDCCA, have limited capability in identifying comprehensive risk genetic loci. There are two reasons. First, all of them demand SNPs to be associated with multi-omics endophenotypes simultaneously, which is overstrict and could lead to a low recall ratio. Second, they ignore the inherent structural information of SNPs such as the LD, and thus are suboptimal for meaningful risk loci identification. ***pc*MTSCCA**

#### The Model

MTSCCA jointly learns multiple SCCA sub-objectives via multi-task learning which could identify comprehensive risk loci for multi-omics problems [22]. However, MTSCCA ignores the multi-omics CEP associations as it only models the relationship between SNPs and multiple types of imaging QTs in parallel. To better make use of multi-omics endophenotypes and their CEP associations, we propose the pairwise endophenotype correlation guided MTSCCA (*pc*MTSCCA). To avoid confusion, the SNP data now is denoted as X, plasma- and CSF-derived proteomic markers and imaging QTs are denoted as Y_1_, Y_2_ and Y_3_ respectively. Then *pc*MTSCCA is defined as follows

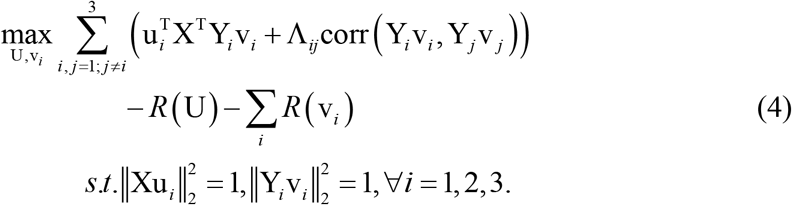

where Λ_*ij*_’s are nonnegative parameters to tune the importance of CEP associations. They can be tuned by the cross-validation strategy. In addition, maximizing Eq. (4) is equivalent to minimizing the following objective

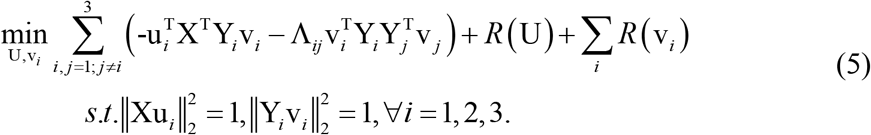

U is the canonical weight for SNPs, and v_1_, v_2_ and v_3_ are those for plasma- and CSF-derived markers and imaging QTs separately. R(U) and R(v_i_)’s are penalisation terms to identify relevant biomarkers. Specifically, R(U) is defined as

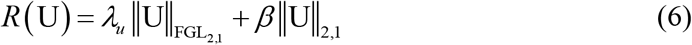

where

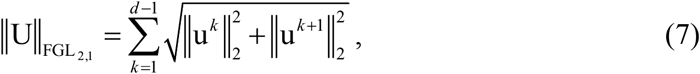

this penalty is the matrix form of FGL [17] and it will reduce to FGL when U degenerates to a vector. This penalty can automatically find out the proximity relationships that extensively exist among SNPs due to the LD in the human genome. λ_*u*_ is a nonnegative parameter that controls the strength of FGL_2,1_ penalty. Additionally, L_2,1_-norm helps identify whether an individual locus affects multi-omics endophenotypes jointly, thereby implicating this locus’ potential pleiotropy. Hence this hybrid penalty is a more reasonable one than that of aforementioned methods in uncovering risk loci. The strength of this L_2,1_-norm is tuned by the nonnegative parameter *β*. In addition, we use L_1_-norm for v_*i*_’s for those multi-omics endophenotypic markers since AD will not attack all endophenotypes of them. Similarly, we use λ_*i*_ (*i* = 1, 2, 3) to control the sparsity of each v_*i*_, and reasonable λ_*i*_’s will help identify those relevant and important endophenotypic markers.

To sum up, *pc*MTSCCA has three advantages. Firstly, it considers both multi-omics endophenotypes and their CEP associations, and thus can be more reasonable than SMCCA, Adaptive SMCCA and RelPMDCCA. Secondly, *pc*MTSCCA has multiple SCCA tasks with each corresponding to the correlation between SNPs and one omics data. On this account, *pc*MTSCCA can make full use of each individual omics data. Thirdly, *pc*MTSCCA employs the novel FGL_2,1_ penalty which takes the LD into consideration while those competitors cannot.

#### Optimization algorithm

The *pc*MTSCCA can be equivalently rewritten as

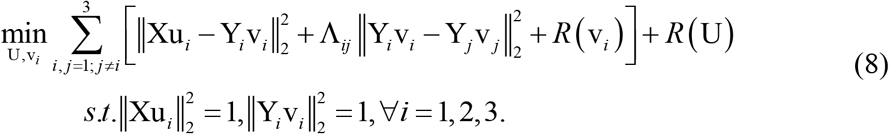

This new formula implies that there is a lower bound. According to [23] (Lemma 2.2), the equality constraints can be finally satisfied by projecting the unconstrained solution onto the L_2_-norm ball. Therefore, we can solve the following unconstrained problem first, *i*.*e*.,

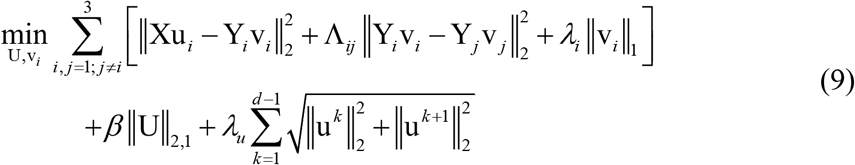

Similar to SMCCA, Eq. (9) is multi-convex in these canonical weights, and we can solve each canonical weight alternatively with those remaining ones fixed. Without loss of generality, we solve the problem with respect to U first. When v_*i*_’s are fixed, the Lagrangian with respect to U is

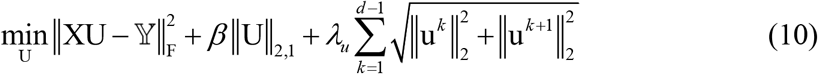

where = [**Y**_1_**v**_1_ **Y**_2_ **v**_2_ **Y**_3_ **v**_3_]. This equation is a multitask regression and can be easily addressed. We take its derivative with respect to U and then set it to zero, i.e.,

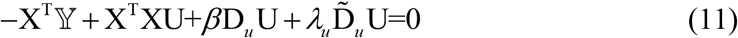

where D_*u*_ is a diagonal matrix, whose diagonal entries are 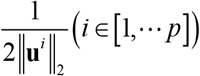, and **D**_*u*_ is another diagonal matrix whose diagonal entries are 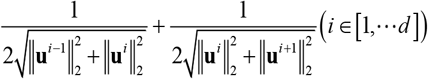 (More details are in [17]). Now we have the solution to Eq. (10) as

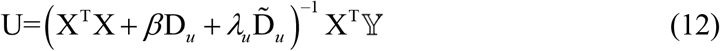

and the solution of U is attained by scaling each u_*i*_, i.e.,

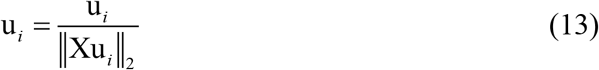

Once obtaining U, we can continue to solve each v_*i*_ alternatively. Similarly, considering the v_*i*_-irrelevant terms as constants in Eq. (9), we take the derivative with respect to v_*i*_, and set it to zero,

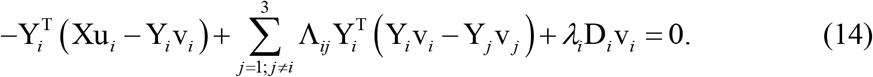

Here D_*i*_ is a diagonal matrix with the *t*-th entry being 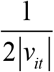, where the range of *t* varies depending on the specific endophenotypic data as we introduced previously. Now we arrive at

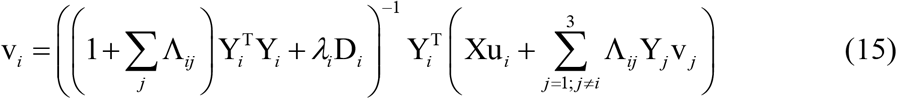

Finally, the solution of v_*i*_ can be attained via the scaling step as follows

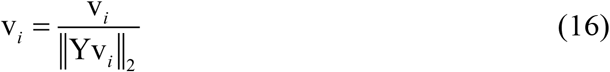

##### Algorithm 1: The *pc*MTSCCA algorithm

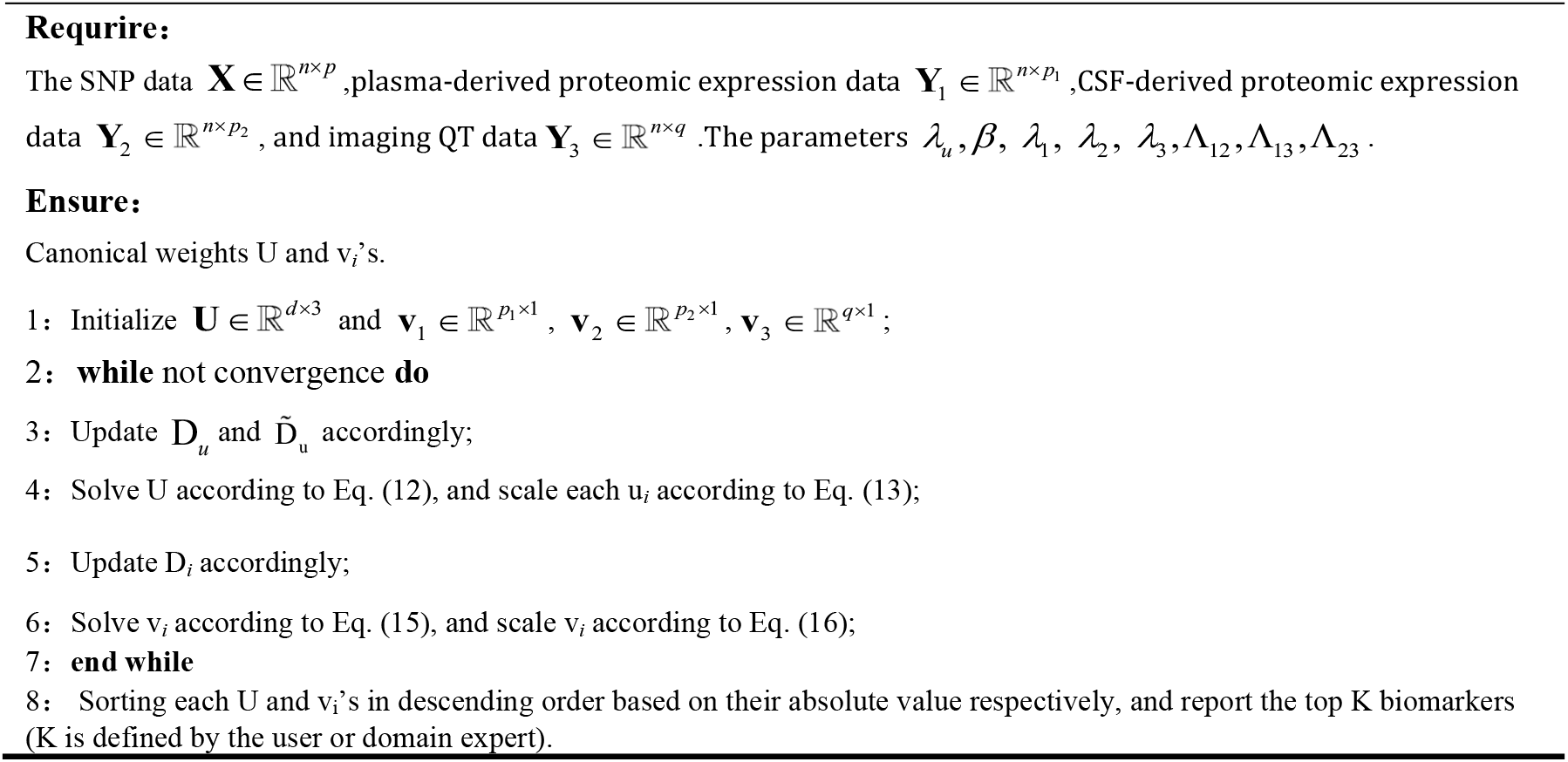

Algorithm 1 contains the pseudo-code of the optimization algorithm. In this algorithm, canonical weights U and v_*i*_’s are alternatively calculated till a pre-defined termination condition is satisfied. Steps 3 and 5 are easy to calculate. In the implementation, we handle Steps 4 and 6 by solving a system of linear equations, which is more efficient than calculating the matrix inverse. Therefore, the proposed algorithm could run with desirable efficiency. According to Eq. (8), the *hoc*MTSCCA objective has the lower bound zero. The iteration algorithm will finally attain a local optimum. In practice, the termination conditions, i.e., max max **U**^*t*+1^ − **U**^*t*^ ≤ *ε* and 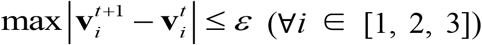, are used to assure efficiency. The tolerance error ϵ was set to 10^−5^ empirically.

#### Extension to high-order endophenotype correlation guided MTSCCA (*hoc*MTSCCA)

Although *pc*MTSCCA is better than existing methods, it only cares about the pairwise correlation between heterogeneous multi-omics endophenotypes. In biomedical studies, the high-order association among multi-omics endophenotypes could also be useful, since the high-order association may implicate novel in-depth clues. To accommodate the high-order CEP association, we propose the high-order endophenotype correlation guided MTSCCA (*hoc*MTSCCA), which is defined as

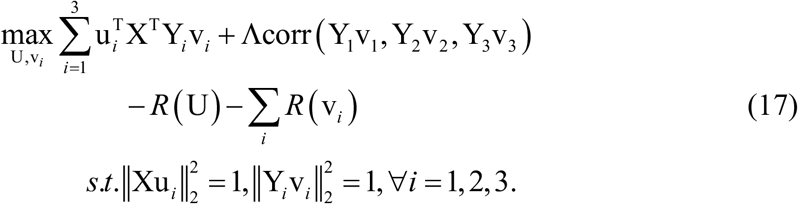

Further, maximizing this equation is equivalent to minimizing the following objective

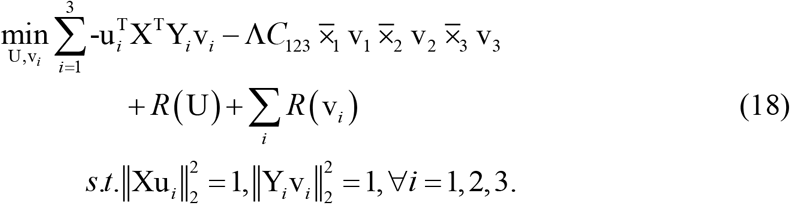

U and v_*i*_’s are canonical weights holding the same meaning to those of *pc*MTSCCA, and so do R(U) and R(v_*i*_)’s. The second term captures the high-order canonical correlation, and Λ is used to control its contribution. According to [24], *C* measures the covariance tensor among multi-omics data which can be calculated from

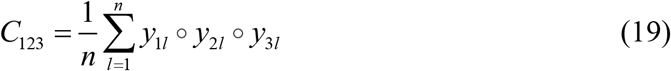

where ° is the tensor product, and 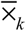 is the k-mode tensor-vector product.

This high-order canonical correlation quantifies the CEP cross multi-omics endophenotypes, and is incorporated into the *hoc*MTSCCA model. On the one hand, *hoc*MTSCCA is similar to *pc*MTSCCA. They both take into account multi-omics endophenotypes and their CEP associations. That is, *hoc*MTSCCA could outperform existing methods too. On the other hand, *hoc*MTSCCA is distinct to *pc*MTSCCA. *hoc*MTSCCA considers the high-order CEP association, while *pc*MTSCCA emphasizes on the pairwise CEP association. This indicates that *hoc*MTSCCA has its unique advantage in identifying risk loci. In a word, *hoc*MTSCCA and *pc*MTSCCA are in the unity of opposites, and both possess better modeling capability than existing multi-omics methods. Since *hoc*MTSCCA follows the same modelling strategy to *pc*MTSCCA, it can be solved in the same way as presented in Algorithm 1, and its convergence is also guaranteed.

## Conclusion

Alzheimer’s disease (AD) is a highly inheritable neurodegenerative brain disorder and many complex traits are observed in AD patients [1]. This inspires us to utilize multiple heterogeneous endophenotypes, derived from multi-omics data, to comprehensively identify genetic loci. Most existing methods cannot make use of both multi-omics endophenotypes and their CEP associations, and thus are suboptimal. Therefore, we proposed two integrated MTSCCA methods, i.e., *pc*MTSCCA and *hoc*MTSCCA, to identify AD-related genetic factors, as well as AD-relevant endophenotypic markers. An efficient optimization algorithm was also proposed.

We compared *pc*MTSCCA and *hoc*MTSCCA with three related multi-omics methods, i.e., SMCCA, Adaptive SMCCA and RelPMDCCA on both simulated and real-world data. The results of synthetic data showed that our methods obtained higher or comparable CCCs and better canonical weights under different circumstances. On the real ADNI data, both *pc*MTSCCA and *hoc*MTSCCA performed better than competitors in terms of CCCs and feature selection. The risk loci, plasma- and CSF derived proteomic markers and imaging QTs identified by our methods were highly related to AD. But comparison methods either yielded too many or too little markers which was undesirable for real biomedical studies. Our methods provided very helpful clues for further in-depth investigation and thus were powerful in multi-omics heterogeneous markers identification. In the future, we will apply both methods to large scales datasets which is important for whole genome sequence analysis.

## Code availability

The inMTSCCA software tool is implemented in MATLAB. The code and manual are available at BioCode of National Genomics Data Center: https://ngdc.cncb.ac.cn/biocode/tools/BT007330. Non-commercial use for academic, government, and non-profit institutions is permitted.

## CRediT author statement

**Du L:** Conceptualization, Methodology, Writing-Original draft preparation. **Zhang J:** Software, Visualization, Formal analysis. **Zhao Y:** Investigation. **Shang M:** Validation. **Guo L:** Writing - Review & Editing. **Han J:** Writing - Review & Editing. All authors have read and approved the final manuscript.

## Acknowledgements

Data collection and sharing for this project was funded by the Alzheimer’s Disease Neuroimaging Initiative (ADNI) (National Institutes of Health Grant U01 AG024904) and DOD ADNI (Department of Defense award number W81XWH-12-2-0012). ADNI is funded by the National Institute on Aging, the National Institute of Biomedical Imaging and Bioengineering, and through generous contributions from the following: AbbVie, Alzheimer’s Association; Alzheimer’s Drug Discovery Foundation; Araclon Biotech; BioClinica, Inc.; Biogen; Bristol-Myers Squibb Company; CereSpir, Inc.; Cogstate; Eisai Inc.; Elan Pharmaceuticals, Inc.; Eli Lilly and Company; EuroImmun; F. HoffmannLa Roche Ltd and its affiliated company Genentech, Inc.; Fujirebio; GE Healthcare; IXICLtd.; Janssen Alzheimer Immunotherapy Research & Development, LLC.; Johnson & Johnson Pharmaceutical Research & Development LLC.;Lumosity; Lundbeck; Merck & Co., Inc.; Meso Scale Diagnostics, LLC.; NeuroRx Research; Neurotrack Technologies; Novartis Pharmaceuticals Corporation; Pfizer Inc.; Piramal Imaging; Servier; Takeda Pharmaceutical Company; and Transition Therapeutics. The Canadian Institutes of Health Research is providing funds to support ADNI clinical sites in Canada. Private sector contributions are facilitated by the Foundation for the National Institutes of Health (www.fnih.org). The grantee organization is the Northern California Institute for Research and Education, and the study is coordinated by the Alzheimer’s Therapeutic Research Institute at the University of Southern California. ADNI data are disseminated by the Laboratory for Neuro Imaging at the University of Southern California.

This work was supported in part by the National Key R&D Program of China [2022ZD0213700]; National Natural Science Foundation of China [61973255, 61936007, 62136004]; Natural Science Basic Research Program of Shaanxi [2020JM-142] and China Postdoctoral Science Foundation [2020T130537] at Northwestern Polytechnical University.

## Tables titles

Table 1 The CCCs (mean ± std.) from 20 times five-fold cross-validation on synthetic data sets.

Table 2 Participant characteristics.

Table 3 Comparison of the CCCs (mean ± std.) from 20 times five-fold cross-validation on ADNI. The best values were shown in bold.

Table 4 Top ten loci of each method based on mean canonical weights.

Table 5 Top ten plasma-derived proteomic markers of each method based on mean canonical weights.

Table 6 Top ten CSF-derived proteomic markers of each method based on mean canonical weights.

Table 7 Top ten brain imaging QTs of each method based on mean canonical weights.

## Supplementary material

**Table S1.**
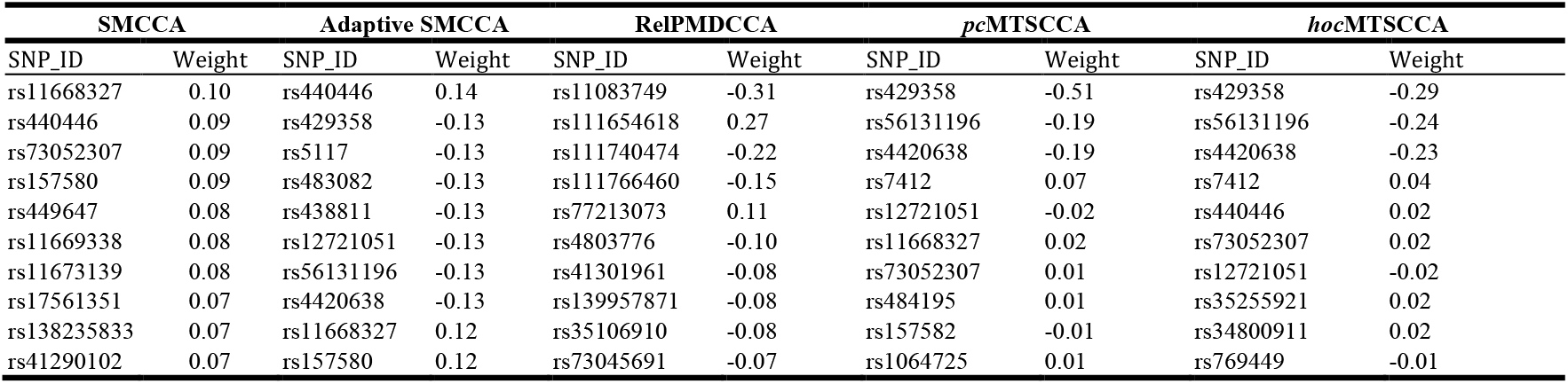
Top ten loci of each method based on mean canonical weights.

**Table S2.**
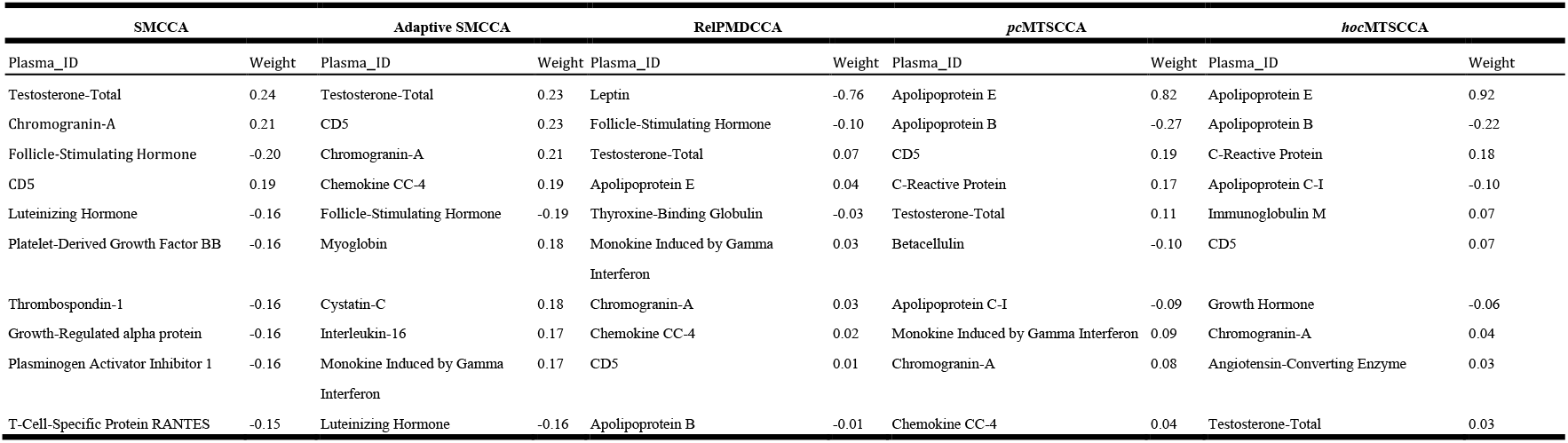
Top ten plasma-derived proteomic markers of each method based on mean canonical weights.

**Table S3.**
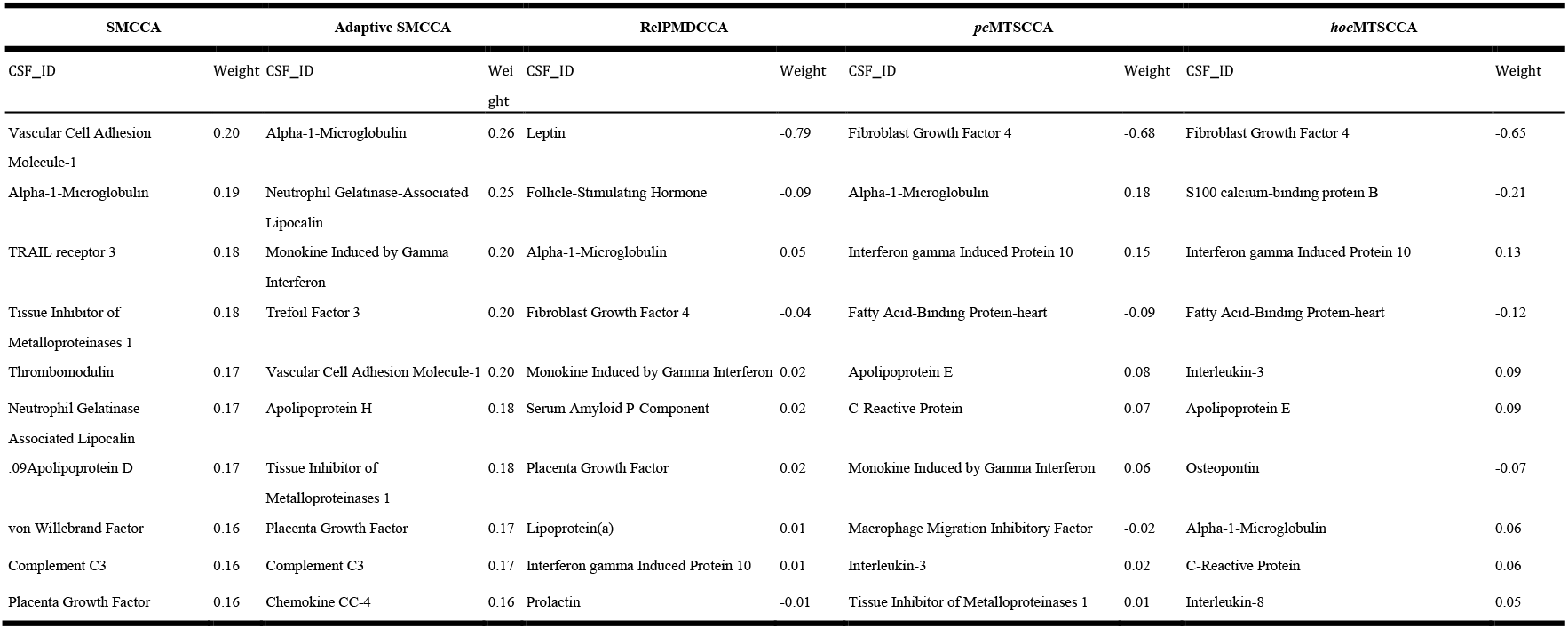
Top ten CSF-derived proteomic markers of each method based on mean canonical weights.

**Table S4.**
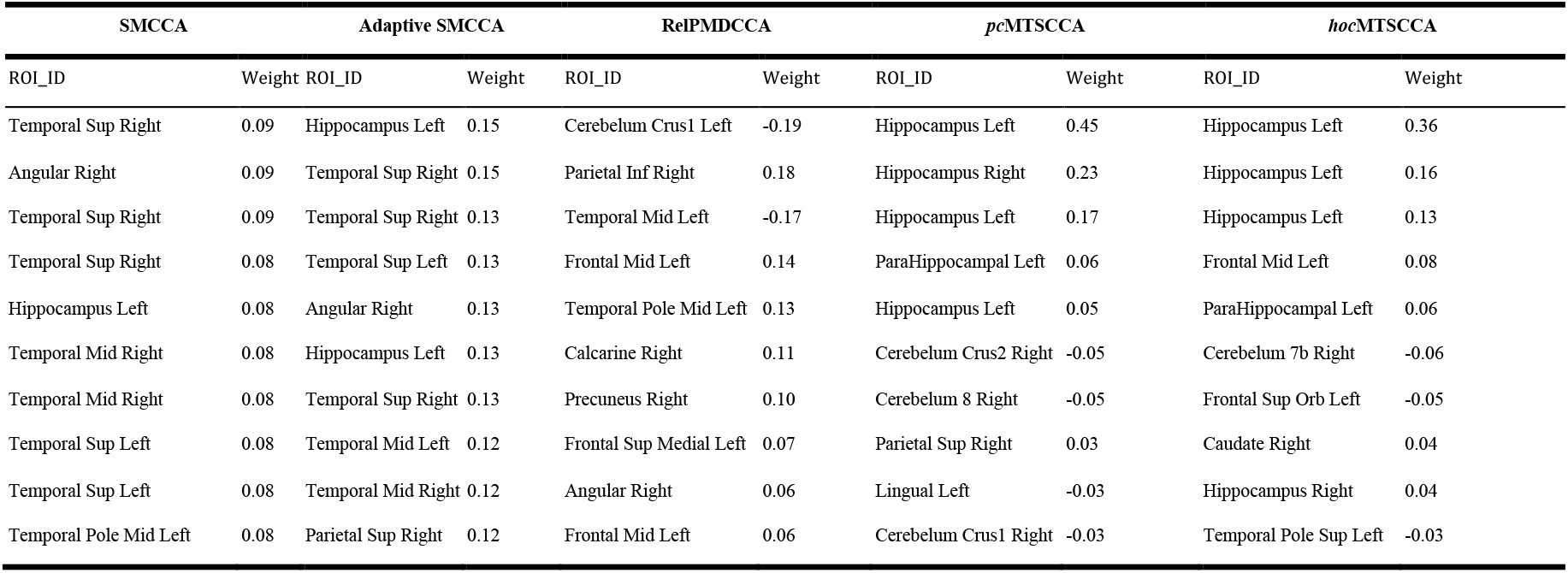
Top ten brain imaging QTs of each method based on mean canonical weights.

