## appendix for "inMTSCCA: An Integrated Multi-task Sparse Canonical Correlation Analysis for Multi-omics Brain Imaging Genetics"

#### Figure legends

**Figure 1 Canonical weights on synthetic data from 20 times five-fold cross-validation**

Each row is a method: (1) Ground truth; (2) SMCCA; (3) Adaptive SMCCA; (4) RelPMDCCA; (5) *pc*MTSCCA; and (6) *hoc*MTSCCA. Each column corresponds to a canonical weight. The first column u is the canonical weight of X, the second column $v_1$ is that of $Y_1$, the third one is the canonical weight of $Y_2$, and the last column is that for $Y_3$. The values in each panel are obtained from 20 times trials. Data source: four simulated data sets with different ground truths. SMCCA, Sparse multiple canonical correlation analysis; RelPMDCCA, Relaxed penalised matrix decomposition CCA; pcMTSCCA, pairwise endophenotype correlation guided multi-task sparse canonical correlation analysis; hocMTSCCA, high-order endophenotype correlation guided MTSCCA

**Figure 2 Canonical weights (mean value) of SNPs from 20 times fivefold cross-validation**

Each row is a method: (1) SMCCA; (2) Adaptive SMCCA; (3) RelPMDCCA; (4) *pc*MTSCCA; and (5) *hoc*MTSCCA. SMCCA, Sparse multiple canonical correlation analysis; RelPMDCCA, Relaxed penalised matrix decomposition CCA; pcMTSCCA, pairwise endophenotype correlation guided multi-task sparse canonical correlation analysis; hocMTSCCA, high-order endophenotype correlation guided MTSCCA; SNP, single nucleotide polymorphism

**Figure 3 Canonical weights (mean value) of plasma-derived proteomic markers from 20 times five-fold cross-validation**

Each row is a method: (1) SMCCA; (2) Adaptive SMCCA; (3) RelPMDCCA; (4) *pc*MTSCCA; and (5) *hoc*MTSCCA. SMCCA, Sparse multiple canonical correlation analysis; RelPMDCCA, Relaxed penalised matrix decomposition CCA; pcMTSCCA, pairwise endophenotype correlation guided multi-task sparse canonical correlation analysis; hocMTSCCA, high-order endophenotype correlation guided MTSCCA

**Figure 4 Canonical weights (mean value) of CSF-derived proteomic markers from 20 times five-fold cross-validation**

Each row is a method: (1) SMCCA; (2) Adaptive SMCCA; (3) RelPMDCCA; (4) *pc*MTSCCA; and (5) *hoc*MTSCCA. SMCCA, Sparse multiple canonical correlation analysis; RelPMDCCA, Relaxed penalised matrix decomposition CCA; pcMTSCCA, pairwise endophenotype correlation guided multi-task sparse canonical correlation analysis; hocMTSCCA, high-order endophenotype correlation guided MTSCCA; CSF, cerebrospinal fluid

**Figure 5 Canonical weights (mean value) of brain imaging QTs from 20 times five-fold cross-validation**

Each row is a method: (1) SMCCA; (2) Adaptive SMCCA; (3) RelPMDCCA; (4) *pc*MTSCCA; and (5) *hoc*MTSCCA. SMCCA, Sparse multiple canonical correlation analysis; RelPMDCCA, Relaxed penalised matrix decomposition CCA; pcMTSCCA, pairwise endophenotype correlation guided multi-task sparse canonical correlation analysis; hocMTSCCA, high-order endophenotype correlation guided MTSCCA; QT, quantitative traits

**Figure 6 *pc*MTSCCA's pairwise correlation**

**A.** The association between identified SNPs and plasma-derived proteomic biomarkers. **B.** The association between identified SNPs and CSF-derived proteomic biomarkers. **C.** The association between identified SNPs and imaging QTs. The '×' symbol indicated that this pairwise association reached the significance level ($p < 0.05$). pcMTSCCA, pairwise endophenotype correlation guided multi-task sparse canonical correlation analysis

**Figure 7 *hoc*MTSCCA's pairwise correlation**

**A.** The association between identified SNPs and plasma-derived proteomic biomarkers. **B.** The association between identified SNPs and CSF-derived proteomic biomarkers. **C.** The association between identified SNPs and imaging QTs. The '×'

symbol indicated that this pairwise association reached the significance level ($p < 0.05$). hocMTSCCA, high-order endophenotype correlation guided MTSCCA

**Figure 8 Pairwise comparisons for SNP and endophenotype among different diagnostic groups such as HC, MCI and AD respectively**

**A.** The concentration of plasma-APOE for different genotypes among three groups.

**B.** The concentration of CSF-APOE for different genotypes among three groups.

**C.** The atrophy in the left hippocampus lobe for different genotypes among three groups.

SNP, single nucleotide polymorphism; HC, healthy control; MCI, mild cognitive impairment; AD, Alzheimer's Disease

**Tables legends**

**Table 1 The CCCs (mean ± std.) from 20 times five-fold cross-validation on synthetic data sets. CCC, canonical correlation coefficient**

**Table 2 Participant characteristics**

**Table 3 Comparison of the CCCs (mean ± std.) from 20 times five-fold cross-validation on ADNI. The best values were shown in bold. CCC, canonical correlation coefficient**

**Table 4 Top ten loci of each method based on mean canonical weights**

**Table 5 Top ten plasma-derived proteomic markers of each method based on mean canonical weights**

**Table 6 Top ten CSF-derived proteomic markers of each method based on mean canonical weights. CSF, cerebrospinal fluid**

**Table 7  Top ten brain imaging QTs of each method based on mean canonical weights. QTs, quantitative traits. QT, quantitative traits**

#### Supplementary material

**Table S1 Top ten loci and their weights of each method based on mean canonical weights**

**Table S2 Top ten plasma-derived proteomic markers and their weights of each method based on mean canonical weights**

**Table S3 Top ten CSF-derived proteomic markers and their weights of each method based on mean canonical weights. CSF, cerebrospinal fluid**

**Table S4 Top ten brain imaging QTs and their weights of each method based on mean canonical weights. QT, quantitative traits**

**Figure 1**

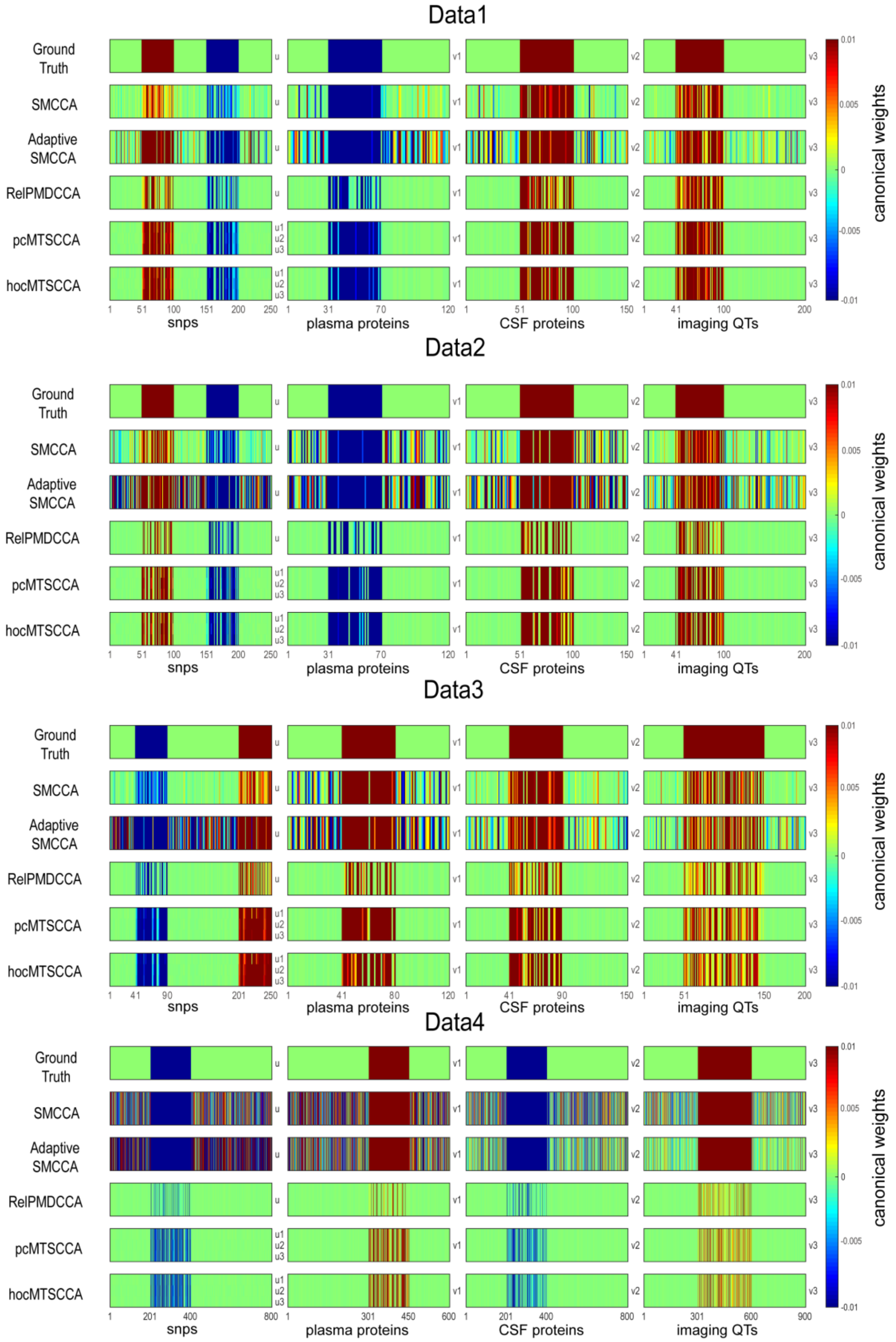
Data1
Data2
Data3
Data4
Ground Truth
SMCCA
Adaptive SMCCA
RelPMDCCA
pcMTSCCA
hocMTSCCA
u
u1
u2
u3
v1
v2
v3
snps
plasma proteins
CSF proteins
imaging QTs
canonical weights
0.01
0.005
0
-0.005
-0.01

### Figure 2

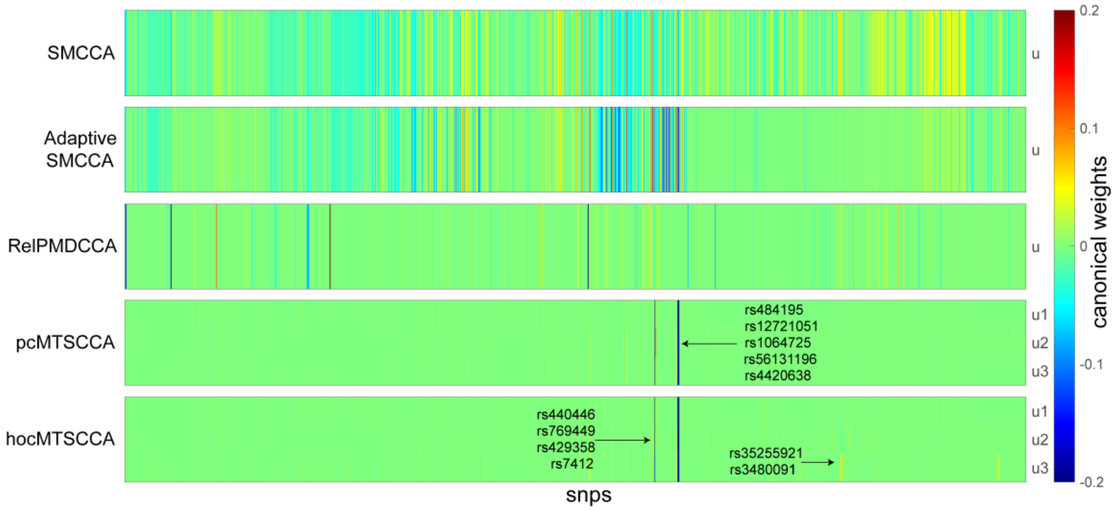


### Figure 3

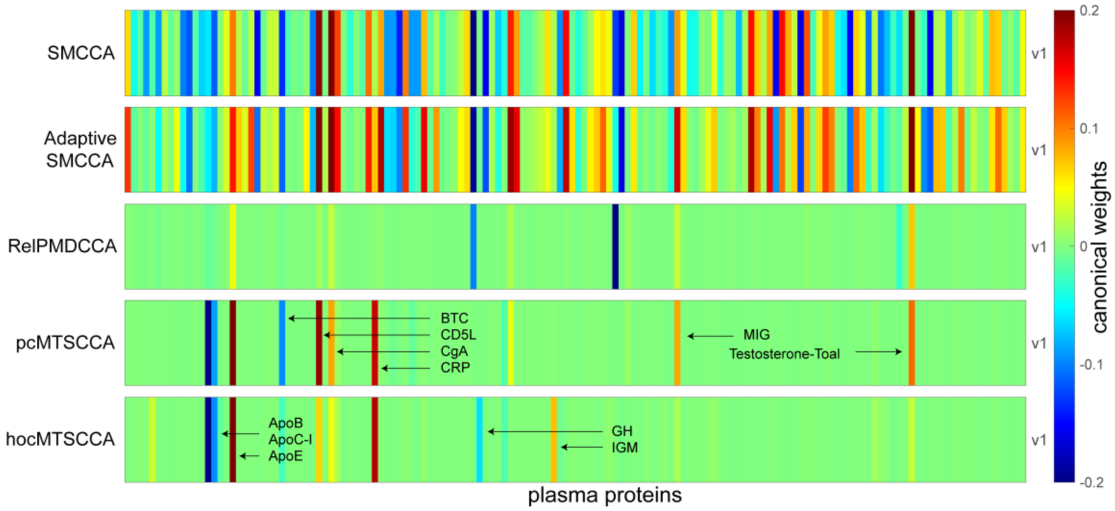


### Figure 4

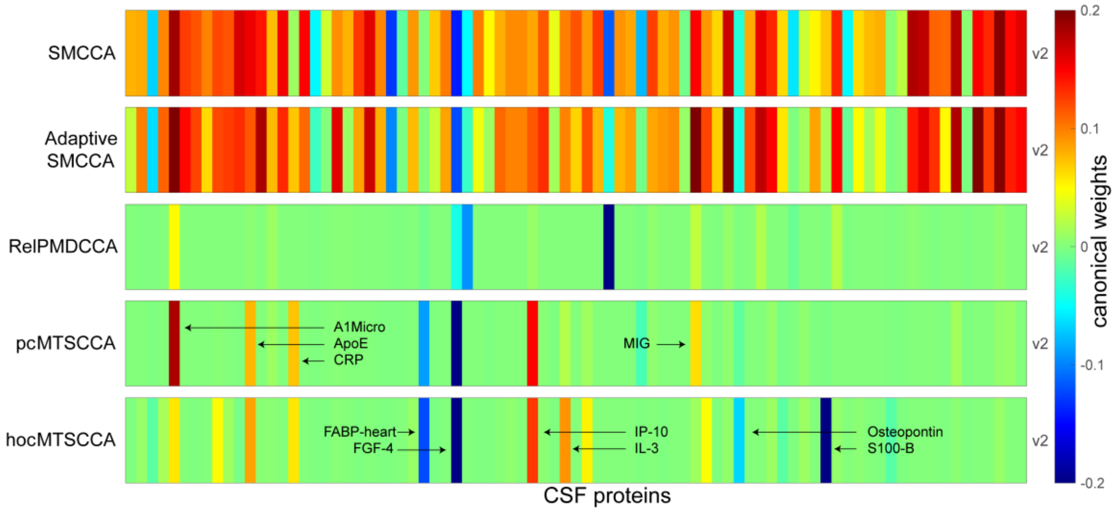

#### Figure 5

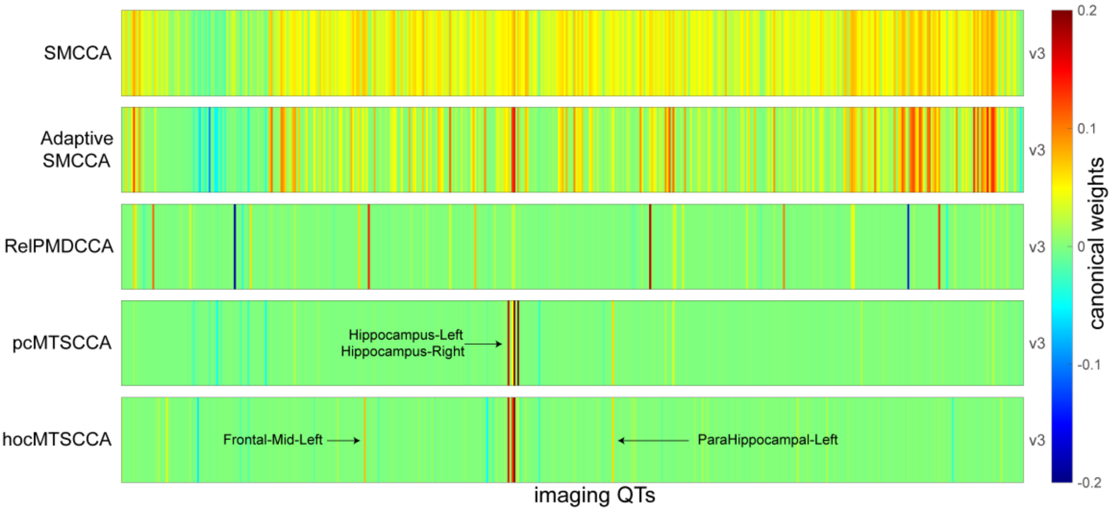


#### Figure 6

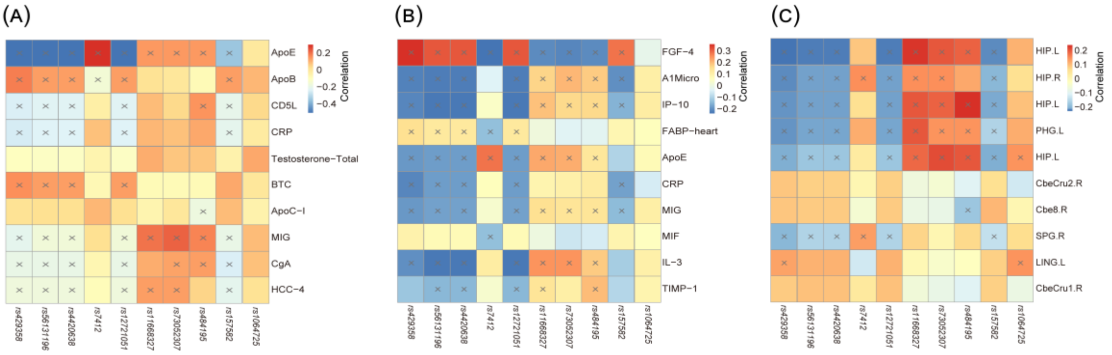


#### Figure 7

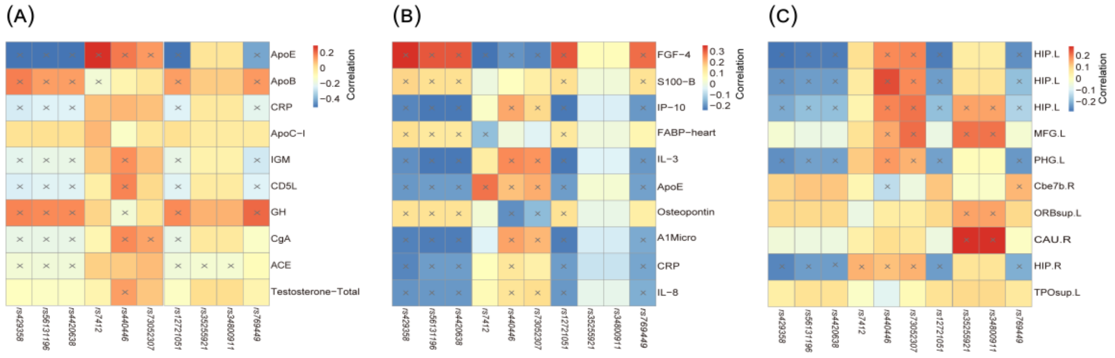

#### Figure 8

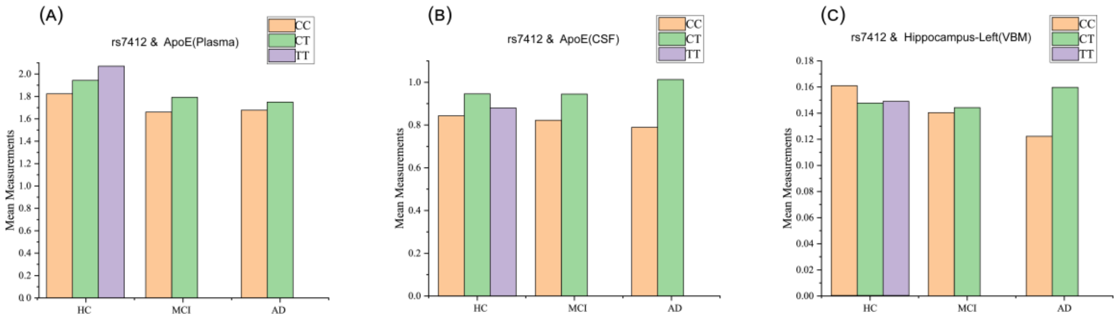


#### Table 1: The CCCs (mean ± std.) from 20 times five-fold cross-validation on synthetic data sets

| | Data1 | | | Data2 | | | Data3 | | | Data4 | | |
|---|---|---|---|---|---|---|---|---|---|---|---|---|
| | **CCC1-1** | **CCC1-2** | **CCC1-3** | **CCC1-1** | **CCC1-2** | **CCC1-3** | **CCC1-1** | **CCC1-2** | **CCC1-3** | **CCC1-1** | **CCC1-2** | **CCC1-3** |
| Training CCCs | | | | | | | | | | | | |
| SMCCA | 0.99 ± 0.00 | 0.99 ± 0.00 | 0.99 ± 0.00 | 0.94 ± 0.01 | 0.94 ± 0.01 | 0.95 ± 0.01 | 0.98 ± 0.00 | 0.99 ± 0.00 | 0.99 ± 0.00 | 0.98 ± 0.00 | 0.98 ± 0.00 | 0.98 ± 0.00 |
| Adaptive SMCCA | 0.99 ± 0.00 | 0.99 ± 0.00 | 0.99 ± 0.00 | 0.96 ± 0.00 | 0.97 ± 0.00 | 0.98 ± 0.01 | 0.98 ± 0.00 | 0.99 ± 0.00 | 0.99 ± 0.00 | 0.98 ± 0.00 | 0.98 ± 0.00 | 0.99 ± 0.00 |
| *Rel*PMDCCA | 0.96 ± 0.00 | 0.97 ± 0.00 | 0.98 ± 0.00 | 0.93 ± 0.01 | 0.93 ± 0.01 | 0.96 ± 0.01 | 0.96 ± 0.01 | 0.98 ± 0.01 | 0.98 ± 0.01 | 0.96 ± 0.01 | 0.96 ± 0.01 | 0.97 ± 0.01 |
| *pc*MTSCCA | 0.99 ± 0.00 | 0.99 ± 0.00 | 0.99 ± 0.00 | 0.96 ± 0.00 | 0.96 ± 0.00 | 0.98 ± 0.00 | 0.98 ± 0.00 | 0.99 ± 0.00 | 0.99 ± 0.00 | 0.98 ± 0.00 | 0.99 ± 0.00 | 0.99 ± 0.00 |
| *hoc*MTSCCA | 0.99 ± 0.00 | 0.99 ± 0.00 | 0.99 ± 0.00 | 0.95 ± 0.00 | 0.96 ± 0.00 | 0.98 ± 0.00 | 0.97 ± 0.00 | 0.99 ± 0.00 | 0.99 ± 0.00 | 0.99 ± 0.00 | 0.98 ± 0.00 | 0.99 ± 0.00 |
| Testing CCCs | | | | | | | | | | | | |
| SMCCA | 0.96 ± 0.00 | 0.97 ± 0.01 | 0.98 ± 0.00 | 0.83 ± 0.05 | 0.80 ± 0.04 | 0.87 ± 0.04 | 0.91 ± 0.03 | 0.96 ± 0.01 | 0.97 ± 0.01 | 0.94 ± 0.01 | 0.94 ± 0.00 | 0.95 ± 0.00 |
| Adaptive SMCCA | 0.96 ± 0.01 | 0.97 ± 0.01 | 0.98 ± 0.00 | 0.83 ± 0.03 | **0.84 ± 0.04** | 0.92 ± 0.03 | 0.91 ± 0.02 | 0.96 ± 0.01 | 0.96 ± 0.01 | 0.94 ± 0.01 | 0.94 ± 0.00 | 0.95 ± 0.00 |
| *Rel*PMDCCA | 0.93 ± 0.03 | 0.94 ± 0.02 | 0.96 ± 0.01 | 0.83 ± 0.05 | 0.82 ± 0.03 | 0.89 ± 0.02 | 0.88 ± 0.04 | 0.94 ± 0.04 | 0.95 ± 0.02 | 0.90 ± 0.02 | 0.89 ± 0.04 | 0.93 ± 0.02 |
| *pc*MTSCCA | **0.97 ± 0.01** | **0.97 ± 0.00** | **0.99 ± 0.00** | **0.84 ± 0.03** | 0.84 ± 0.05 | **0.93 ± 0.02** | **0.92 ± 0.02** | **0.98 ± 0.01** | **0.98 ± 0.01** | **0.95 ± 0.02** | 0.95 ± 0.02 | **0.96 ± 0.01** |
| *hoc*MTSCCA | **0.97 ± 0.01** | 0.97 ± 0.01 | **0.99 ± 0.00** | 0.83 ± 0.03 | **0.84 ± 0.04** | **0.93 ± 0.02** | 0.91 ± 0.02 | 0.97 ± 0.02 | **0.98 ± 0.01** | **0.95 ± 0.02** | **0.95 ± 0.01** | 0.95 ± 0.02 |

*Note*: CCC, Canonical correlation coefficient. The CCC between X and $Y_1$ as CCC1-1, that between X and $Y_2$ as CCC1-2 and so on. Bold fonts represent the best results for the method. SMCCA, Sparse multiple canonical correlation analysis; RelPMDCCA, Relaxed penalised matrix decomposition CCA; pcMTSCCA, pairwise endophenotype correlation guided multi-task sparse canonical correlation analysis; hocMTSCCA, high-order endophenotype correlation guided MTSCCA.

#### Table 2: Participant characteristics

| | HC | MCI | AD |
|---|---|---|---|
| Num | 42 | 137 | 65 |
| Gender (M/F, %) | 52.38/47.62 | 69.34/30.66 | 55.38/44.62 |
| Handedness (R/L, %) | 90.48/9.52 | 92.70/7.30 | 98.46/1.54 |
| Age (mean ± std) | 75.40±5.80 | 74.13±7.22 | 74.75±7.67 |
| Education (mean ± std) | 15.88±2.77 | 16.03±2.98 | 15.12±3.05 |

*Note*: HC, healthy control; MCI, mild cognitive impairment; AD, Alzheimer's Disease.

**Table 3: Comparison of the CCCs (mean ± std.) from 20 times five-fold cross-validation on ADNI. The best values were shown in bold**

| | Training CCCs | | |
|---|---|---|---|
| | SNP-Plasma | SNP-CSF | SNP-VBM |
| SMCCA | 0.33 ± 0.04 | 0.33 ± 0.04 | 0.29 ± 0.02 |
| Adaptive SMCCA | 0.36 ± 0.03 | 0.34 ± 0.03 | 0.23 ± 0.02 |
| *Rel*PMDCCA | 0.44 ± 0.05 | 0.46 ± 0.03 | **0.50 ± 0.04** |
| *pc*MTSCCA | 0.66 ± 0.04 | 0.44 ± 0.04 | 0.37 ± 0.04 |
| *hoc*MTSCCA | **0.66 ± 0.03** | **0.50 ± 0.04** | 0.44 ± 0.04 |

| | Testing CCCs | | |
|---|---|---|---|
| | SNP-Plasma | SNP-CSF | SNP-VBM |
| SMCCA | 0.13 ± 0.09 | 0.18 ± 0.10 | 0.09 ± 0.06 |
| Adaptive SMCCA | 0.17 ± 0.11 | 0.21 ± 0.11 | 0.10 ± 0.07 |
| *Rel*PMDCCA | 0.13 ± 0.12 | 0.11 ± 0.08 | 0.12 ± 0.09 |
| *pc*MTSCCA | **0.56 ± 0.11** | 0.30 ± 0.12 | **0.17 ± 0.11** |
| *hoc*MTSCCA | **0.56 ± 0.11** | **0.31 ± 0.13** | 0.15 ± 0.10 |

*Note*: SMCCA, Sparse multiple canonical correlation analysis; RelPMDCCA, Relaxed penalised matrix decomposition CCA; pcMTSCCA, pairwise endophenotype correlation guided multi-task sparse canonical correlation analysis; hocMTSCCA, high-order endophenotype correlation guided MTSCCA; CCC, Canonical correlation coefficient.

**Table 4: Top ten loci of each method based on mean canonical weights**

| SMCCA | Adaptive SMCCA | *Rel*PMDCCA | *pc*MTSCCA | *hoc*MTSCCA |
|---|---|---|---|---|
| rs11668327 | rs440446 | rs11083749 | rs429358 | rs429358 |
| rs440446 | rs429358 | rs111654618 | rs56131196 | rs56131196 |
| rs73052307 | rs5117 | rs111740474 | rs4420638 | rs4420638 |
| rs157580 | rs483082 | rs111766460 | rs7412 | rs7412 |
| rs449647 | rs438811 | rs77213073 | rs12721051 | rs440446 |
| rs11669338 | rs12721051 | rs4803776 | rs11668327 | rs73052307 |
| rs11673139 | rs56131196 | rs41301961 | rs73052307 | rs12721051 |
| rs17561351 | rs4420638 | rs139957871 | rs484195 | rs35255921 |
| rs138235833 | rs11668327 | rs35106910 | rs157582 | rs34800911 |
| rs41290102 | rs157580 | rs73045691 | rs1064725 | rs769449 |

*Note*: SMCCA, Sparse multiple canonical correlation analysis; RelPMDCCA, Relaxed penalised matrix decomposition CCA; pcMTSCCA, pairwise endophenotype correlation guided multi-task sparse canonical correlation analysis; hocMTSCCA, high-order endophenotype correlation guided MTSCCA.

**Table 5: Top ten plasma-derived proteomic markers of each method based on mean canonical weights**

| SMCCA | Adaptive SMCCA | *Rel*PMDCCA | *pc*MTSCCA | *hoc*MTSCCA |
|---|---|---|---|---|
| Testosterone | Testosterone | Leptin | ApoE | ApoE |
| CgA | CD5L | FSH | ApoB | ApoB |
| FSH | CgA | Testosterone | CD5L | CRP |
| CD5L | HCC-4 | ApoE | CRP | ApoC-I |
| LH | FSH | TBG | Testosterone | IGM |
| PDGF-BB | Myoglobin | MIG | BTC | CD5L |
| Thrombospondin-1 | Cystatin-C | CgA | ApoC-I | GH |
| GRO-alpha | IL-16 | HCC-4 | MIG | CgA |
| PAI-1 | MIG | CD5L | CgA | ACE |
| RANTES | LH | ApoB | HCC-4 | Testosterone |

*Note*: SMCCA, Sparse multiple canonical correlation analysis; RelPMDCCA, Relaxed penalised matrix decomposition CCA; pcMTSCCA, pairwise endophenotype correlation guided multi-task sparse canonical correlation analysis; hocMTSCCA, high-order endophenotype correlation guided MTSCCA.

**Table 6: Top ten CSF-derived proteomic markers of each method based on mean canonical weights**

| **SMCCA** | **Adaptive SMCCA** | ***Rel*PMDCCA** | ***pc*MTSCCA** | ***hoc*MTSCCA** |
|---|---|---|---|---|
| VCAM-1 | A1Micro | Leptin | FGF-4 | FGF-4 |
| A1Micro | NGAL | FSH | A1Micro | S100-B |
| TRAIL-R3 | MIG | A1Micro | IP-10 | IP-10 |
| TIMP-1 | TFF3 | FGF-4 | FABP-heart | FABP-heart |
| TM | VCAM-1 | MIG | ApoE | IL-3 |
| NGAL | ApoH | SAP | CRP | ApoE |
| ApoD | TIMP-1 | PLGF | MIG | Osteopontin |
| vWF | PLGF | LPa | MIF | A1Micro |
| C3 | C3 | IP-10 | IL-3 | CRP |
| PLGF | HCC-4 | PRL | TIMP-1 | IL-8 |

*Note*: SMCCA, Sparse multiple canonical correlation analysis; RelPMDCCA, Relaxed penalised matrix decomposition CCA; pcMTSCCA, pairwise endophenotype correlation guided multi-task sparse canonical correlation analysis; hocMTSCCA, high-order endophenotype correlation guided MTSCCA; CSF, cerebrospinal fluid.

**Table 7: Top ten brain imaging QTs of each method based on mean canonical weights**

| **SMCCA** | **Adaptive SMCCA** | ***Rel*PMDCCA** | ***pc*MTSCCA** | ***hoc*MTSCCA** |
|---|---|---|---|---|
| STG.R | HIP.L | CbeCru1.L | HIP.L | HIP.L |
| ANG.R | STG.R | IPL.R | HIP.R | HIP.L |
| STG.R | STG.R | MTG.L | HIP.L | HIP.L |
| STG.R | STG.L | MFG.L | PHG.L | MFG.L |
| HIP.L | ANG.R | TPOmid.L | HIP.L | PHG.L |
| MTG.R | HIP.L | CAL.R | CbeCru2.R | Cbe7b.R |
| MTG.R | STG.R | PCUN.R | Cbe8.R | ORBsup.L |
| STG.L | MTG.L | SFGmed.L | SPG.R | CAU.R |
| STG.L | MTG.R | ANG.R | LING.L | HIP.R |
| TPOmid.L | SPG.R | MFG.L | CbeCru1.R | TPOsup.L |

*Note*: SMCCA, Sparse multiple canonical correlation analysis; RelPMDCCA, Relaxed penalised matrix decomposition CCA; pcMTSCCA, pairwise endophenotype correlation guided multi-task sparse canonical correlation analysis; hocMTSCCA, high-order endophenotype correlation guided MTSCCA; QT, quantitative traits.

### Supplementary material

#### Table S1 Top ten loci and their weights of each method based on mean canonical weights

| SMCCA | | Adaptive SMCCA | | RelPMDCCA | | *pc*MTSCCA | | *hoc*MTSCCA | |
|---|---|---|---|---|---|---|---|---|---|
| SNP_ID | Weight | SNP_ID | Weight | SNP_ID | Weight | SNP_ID | Weight | SNP_ID | Weight |
| rs11668327 | 0.10 | rs440446 | 0.14 | rs11083749 | -0.31 | rs429358 | -0.51 | rs429358 | -0.29 |
| rs440446 | 0.09 | rs429358 | -0.13 | rs111654618 | 0.27 | rs56131196 | -0.19 | rs56131196 | -0.24 |
| rs73052307 | 0.09 | rs5117 | -0.13 | rs111740474 | -0.22 | rs4420638 | -0.19 | rs4420638 | -0.23 |
| rs157580 | 0.09 | rs483082 | -0.13 | rs111766460 | -0.15 | rs7412 | 0.07 | rs7412 | 0.04 |
| rs449647 | 0.08 | rs438811 | -0.13 | rs77213073 | 0.11 | rs12721051 | -0.02 | rs440446 | 0.02 |
| rs11669338 | 0.08 | rs12721051 | -0.13 | rs4803776 | -0.10 | rs11668327 | 0.02 | rs73052307 | 0.02 |
| rs11673139 | 0.08 | rs56131196 | -0.13 | rs41301961 | -0.08 | rs73052307 | 0.01 | rs12721051 | -0.02 |
| rs17561351 | 0.07 | rs4420638 | -0.13 | rs139957871 | -0.08 | rs484195 | 0.01 | rs35255921 | 0.02 |
| rs138235833 | 0.07 | rs11668327 | 0.12 | rs35106910 | -0.08 | rs157582 | -0.01 | rs34800911 | 0.02 |
| rs41290102 | 0.07 | rs157580 | 0.12 | rs73045691 | -0.07 | rs1064725 | 0.01 | rs769449 | -0.01 |

*Note*: SMCCA, Sparse multiple canonical correlation analysis; RelPMDCCA, Relaxed penalised matrix decomposition CCA; pcMTSCCA, pairwise endophenotype correlation guided multi-task sparse canonical correlation analysis; hocMTSCCA, high-order endophenotype correlation guided MTSCCA.

#### Table S2 Top ten plasma-derived proteomic markers and their weights of each method based on mean canonical weights

| SMCCA | | Adaptive SMCCA | | RelPMDCCA | | *pc*MTSCCA | | *hoc*MTSCCA | |
|---|---|---|---|---|---|---|---|---|---|
| Plasma_ID | Weight | Plasma_ID | Weight | Plasma_ID | Weight | Plasma_ID | Weight | Plasma_ID | Weight |
| Testosterone-Total | 0.24 | Testosterone-Total | 0.23 | Leptin | -0.76 | Apolipoprotein E | 0.82 | Apolipoprotein E | 0.92 |
| Chromogranin-A | 0.21 | CD5 | 0.23 | Follicle-Stimulating Hormone | -0.10 | Apolipoprotein B | -0.27 | Apolipoprotein B | -0.22 |
| Follicle-Stimulating Hormone | -0.20 | Chromogranin-A | 0.21 | Testosterone-Total | 0.07 | CD5 | 0.19 | C-Reactive Protein | 0.18 |
| CD5 | 0.19 | Chemokine CC-4 | 0.19 | Apolipoprotein E | 0.04 | C-Reactive Protein | 0.17 | Apolipoprotein C-I | -0.10 |
| Luteinizing Hormone | -0.16 | Follicle-Stimulating Hormone | -0.19 | Thyroxine-Binding Globulin | -0.03 | Testosterone-Total | 0.11 | Immunoglobulin M | 0.07 |
| Platelet-Derived Growth Factor BB | -0.16 | Myoglobin | 0.18 | Monokine Induced by Gamma Interferon | 0.03 | Betacellulin | -0.10 | CD5 | 0.07 |
| Thrombospondin-1 | -0.16 | Cystatin-C | 0.18 | Chromogranin-A | 0.03 | Apolipoprotein C-I | -0.09 | Growth Hormone | -0.06 |
| Growth-Regulated alpha protein | -0.16 | Interleukin-16 | 0.17 | Chemokine CC-4 | 0.02 | Monokine Induced by Gamma Interferon | 0.09 | Chromogranin-A | 0.04 |
| Plasminogen Activator Inhibitor 1 | -0.16 | Monokine Induced by Gamma Interferon | 0.17 | CD5 | 0.01 | Chromogranin-A | 0.08 | Angiotensin-Converting Enzyme | 0.03 |
| T-Cell-Specific Protein RANTES | -0.15 | Luteinizing Hormone | -0.16 | Apolipoprotein B | -0.01 | Chemokine CC-4 | 0.04 | Testosterone-Total | 0.03 |

*Note*: SMCCA, Sparse multiple canonical correlation analysis; RelPMDCCA, Relaxed penalised matrix decomposition CCA; pcMTSCCA, pairwise endophenotype correlation guided multi-task sparse canonical correlation analysis; hocMTSCCA, high-order endophenotype correlation guided MTSCCA.

#### Table S3 Top ten CSF-derived proteomic markers and their weights of each method based on mean canonical weights

| SMCCA | | Adaptive SMCCA | | RelPMDCCA | | *pc*MTSCCA | | *hoc*MTSCCA | |
|---|---|---|---|---|---|---|---|---|---|
| CSF_ID | Weight | CSF_ID | Weight | CSF_ID | Weight | CSF_ID | Weight | CSF_ID | Weight |
| Vascular Cell Adhesion Molecule-1 | 0.20 | Alpha-1-Microglobulin | 0.26 | Leptin | -0.79 | Fibroblast Growth Factor 4 | -0.68 | Fibroblast Growth Factor 4 | -0.65 |
| Alpha-1-Microglobulin | 0.19 | Neutrophil Gelatinase-Associated Lipocalin | 0.25 | Follicle-Stimulating Hormone | -0.09 | Alpha-1-Microglobulin | 0.18 | S100 calcium-binding protein B | -0.21 |
| TRAIL receptor 3 | 0.18 | Monokine Induced by Gamma Interferon | 0.20 | Alpha-1-Microglobulin | 0.05 | Interferon gamma Induced Protein 10 | 0.15 | Interferon gamma Induced Protein 10 | 0.13 |
| Tissue Inhibitor of Metalloproteinases 1 | 0.18 | Trefoil Factor 3 | 0.20 | Fibroblast Growth Factor 4 | -0.04 | Fatty Acid-Binding Protein-heart | -0.09 | Fatty Acid-Binding Protein-heart | -0.12 |
| Thrombomodulin | 0.17 | Vascular Cell Adhesion Molecule-1 | 0.20 | Monokine Induced by Gamma Interferon | 0.02 | Apolipoprotein E | 0.08 | Interleukin-3 | 0.09 |
| Neutrophil Gelatinase-Associated Lipocalin | 0.17 | Apolipoprotein H | 0.18 | Serum Amyloid P-Component | 0.02 | C-Reactive Protein | 0.07 | Apolipoprotein E | 0.09 |
| .09Apolipoprotein D | 0.17 | Tissue Inhibitor of Metalloproteinases 1 | 0.18 | Placenta Growth Factor | 0.02 | Monokine Induced by Gamma Interferon | 0.06 | Osteopontin | -0.07 |
| von Willebrand Factor | 0.16 | Placenta Growth Factor | 0.17 | Lipoprotein(a) | 0.01 | Macrophage Migration Inhibitory Factor | -0.02 | Alpha-1-Microglobulin | 0.06 |
| Complement C3 | 0.16 | Complement C3 | 0.17 | Interferon gamma Induced Protein 10 | 0.01 | Interleukin-3 | 0.02 | C-Reactive Protein | 0.06 |
| Placenta Growth Factor | 0.16 | Chemokine CC-4 | 0.16 | Prolactin | -0.01 | Tissue Inhibitor of Metalloproteinases 1 | 0.01 | Interleukin-8 | 0.05 |

*Note*: SMCCA, Sparse multiple canonical correlation analysis; RelPMDCCA, Relaxed penalised matrix decomposition CCA; pcMTSCCA, pairwise endophenotype correlation guided multi-task sparse canonical correlation analysis; hocMTSCCA, high-order endophenotype correlation guided MTSCCA; CSF, cerebrospinal fluid.

#### Table S4 Top ten brain imaging QTs and their weights of each method based on mean canonical weights

| SMCCA | | Adaptive SMCCA | | RelPMDCCA | | *pc*MTSCCA | | *hoc*MTSCCA | |
|---|---|---|---|---|---|---|---|---|---|
| ROI_ID | Weight | ROI_ID | Weight | ROI_ID | Weight | ROI_ID | Weight | ROI_ID | Weight |
| Temporal Sup Right | 0.09 | Hippocampus Left | 0.15 | Cerebelum Crus1 Left | -0.19 | Hippocampus Left | 0.45 | Hippocampus Left | 0.36 |
| Angular Right | 0.09 | Temporal Sup Right | 0.15 | Parietal Inf Right | 0.18 | Hippocampus Right | 0.23 | Hippocampus Left | 0.16 |
| Temporal Sup Right | 0.09 | Temporal Sup Right | 0.13 | Temporal Mid Left | -0.17 | Hippocampus Left | 0.17 | Hippocampus Left | 0.13 |
| Temporal Sup Right | 0.08 | Temporal Sup Left | 0.13 | Frontal Mid Left | 0.14 | ParaHippocampal Left | 0.06 | Frontal Mid Left | 0.08 |
| Hippocampus Left | 0.08 | Angular Right | 0.13 | Temporal Pole Mid Left | 0.13 | Hippocampus Left | 0.05 | ParaHippocampal Left | 0.06 |
| Temporal Mid Right | 0.08 | Hippocampus Left | 0.13 | Calcarine Right | 0.11 | Cerebelum Crus2 Right | -0.05 | Cerebelum 7b Right | -0.06 |
| Temporal Mid Right | 0.08 | Temporal Sup Right | 0.13 | Precuneus Right | 0.10 | Cerebelum 8 Right | -0.05 | Frontal Sup Orb Left | -0.05 |
| Temporal Sup Left | 0.08 | Temporal Mid Left | 0.12 | Frontal Sup Medial Left | 0.07 | Parietal Sup Right | 0.03 | Caudate Right | 0.04 |
| Temporal Sup Left | 0.08 | Temporal Mid Right | 0.12 | Angular Right | 0.06 | Lingual Left | -0.03 | Hippocampus Right | 0.04 |
| Temporal Pole Mid Left | 0.08 | Parietal Sup Right | 0.12 | Frontal Mid Left | 0.06 | Cerebelum Crus1 Right | -0.03 | Temporal Pole Sup Left | -0.03 |

*Note*: SMCCA, Sparse multiple canonical correlation analysis; RelPMDCCA, Relaxed penalised matrix decomposition CCA; pcMTSCCA, pairwise endophenotype correlation guided multi-task sparse canonical correlation analysis; hocMTSCCA, high-order endophenotype correlation guided MTSCCA; QT, quantitative traits.
